# Inferring ligand-receptor cellular networks from bulk and spatial transcriptomic datasets with BulkSignalR

**DOI:** 10.1101/2022.11.17.516911

**Authors:** Jean-Philippe Villemin, Laia Bassaganyas, Didier Pourquier, Florence Boissiere, Simon Cabello-Aguilar, Evelyne Crapez, Rita Tanos, Emmanuel Cornillot, Andrei Turtoi, Jacques Colinge

**Affiliations:** Institut de Recherche en Cancérologie de Montpellier (IRCM), Inserm U 1194, Montpellier, France; Université de Montpellier, Montpellier, France; Institut régional du Cancer Montpellier (ICM), Montpellier, France; Faculté de Pharmacie, Université de Montpellier, Montpellier, France; Faculté de Médecine, Université de Montpellier, Montpellier, France

## Abstract

The study of cellular networks mediated by ligand-receptor interactions has attracted much attention recently owing to single-cell omics. However, rich collections of bulk data accompanied with clinical information exists and continue to be generated with no equivalent in single-cell so far. In parallel, spatial transcriptomic (ST) analyses represent a revolutionary tool in biology. A large number of ST projects rely on multicellular resolution, for instance the Visium^™^ platform, where several cells are analyzed at each location, thus producing localized bulk data. Here, we describe BulkSignalR, a R package to infer ligand-receptor networks from bulk data. BulkSignalR integrates ligand-receptor interactions with downstream pathways to estimate statistical significance. A range of visualization methods complement the statistics, including functions dedicated to spatial data. We demonstrate BulkSignalR relevance using different bulk datasets, including new Visium liver metastasis ST data, with experimental validation of selected interactions. A comparison with other ST packages shows the significantly higher quality of BulkSignalR inferences. BulkSignalR can be applied to any species thanks to its built-in generic ortholog mapping functionality.

## INTRODUCTION

The dialog of cells in a tissue through the secretion of ligands and sensing by receptors plays an essential role in development, homeostasis, and diseases (1). The advent of single-cell omics has led to remarkable progresses in the analysis of the cell composition and ligand-receptor networks within a tissue (2–5). Nevertheless, these technologies remain expensive and single-cell data on cohorts are limited compared with bulk datasets, particularly for patient cohorts associated with clinical data. Moreover, although bulk techniques cannot give direct insights into the activity/function of individual cell types, they are more sensitive for detecting low-abundance molecules. Therefore, tools to untangle cellular networks from bulk data are needed as a complementary solution to single-cell studies.

Here, we describe BulkSignalR, a R package that builds on our previous work on bulk (6) and single-cell data (7). BulkSignalR exploits reference databases of known ligand-receptor interactions (LRIs), gene or protein interactions, and biological pathways to assess the significance of correlation patterns between a ligand, its putative receptor, and the targets of the downstream pathway. This integrated modeling provides the increased specificity that is required by the convoluted bulk format. It also allows generating gene signatures to report the LRI activity and their downstream consequences. This may facilitate sample comparison and may be used for patient stratification. As BulkSignalR uses correlation patterns to determine the statistical significance of LRIs, datasets can be analyzed without any prior knowledge of sample groups or clusters.

Despite the popularity of LRI inference in the single-cell bioinformatics community with many existing tools, its equivalent in bulk data has not attracted much attention. A seminal paper explored LRIs in non-small-cell lung cancer (NSCLC) (8) using an empirical algorithm, called CCCExplorer, that requires separate bulk transcriptome datasets for different cell populations. Exploiting dual mouse and human microarrays, *i.e*., dual transcriptomes, Komurov (9) proposed an algorithm to infer interactions between cancer and stromal cells using bulk data from a lung adenocarcinoma mouse xenografts model. Among the studies on LRI inference from single-cell data, two tools indicate in their documentation that they may be applied to bulk data. CellPhoneDB (10) mentions the possibility to process bulk data provided they were obtained from pure cell populations, and ICELLNET (11) is able to exploit two separated bulk datasets to predict interactions. Due to the need for separated bulk datasets, CCCExplorer, CellPhoneDB, and ICELLNET cannot be compared with BulkSignalR directly. They offer a less general approach.

After single-cell omics, spatial transcriptomics (ST) (12) is another revolution in functional genomics. Spatial data are often obtained at multicellular resolution, *e.g*., with the popular Visium^™^ system, and therefore, they are localized bulk data. Consequently, bulk-specific approaches could be used to assess ST data analysis. However, most software tools developed for ST target single-cell or subcellular resolution datasets, where the individual transcriptomes of adjacent or nearby cells can be directly accessed. By simply adjusting few parameters, we found that BulkSignalR could be used for multicellular resolution spatial analyses successfully. We then compared the performance of BulkSignalR and of three tools for multicellular resolution analyses, CellPhoneDB (10), stLearn (13), and SpaTalk (14).

Lastly, we generated an original Visium^™^ dataset from four colorectal cancer liver metastases (CRC-LM) to validate a selection of BulkSignalR predictions by immunofluorescence (IF) analysis.

## MATERIALS AND METHODS

### Expression data and randomized expression data

We denote with *A* the *n*×*m* matrix that represents the expression of *n* genes (or proteins) in *m* samples. To compute the null distributions of the Spearman rank correlation coefficients, we need to generate randomized expression datasets. To do this, we assign each gene to *b* equally sized bins of comparable average (over the samples) expression levels and we shuffle genes within the same bin. By default, we use *b* = 20.

### Ligand-receptor database and pathways

We import known LRIs (the LR*db* database), pathways, and a global reference gene/protein interaction network from SingleCellSignalR (7). For each receptor in a pathway, we identify genes the expression of which might correlate with the receptor expression (see Results), and call them target genes.

### Null distributions of the Spearman rank correlation coefficients

BulkSignalR statistical model (below) requires null distributions of Spearman correlation coefficients between a ligand and a receptor (L-R null distributions) and also between a receptor and the genes involved in a pathway that includes that receptor. We called these genes target genes and the corresponding null distribution is the R-T null distribution. By default, an automated algorithm selects the appropriate statistical model for these null distributions because their shapes depend on the dataset.

Empirical random Spearman L-R correlations are obtained by generating *r_1_*. randomized expression datasets. For each dataset, Spearman correlation (across samples) is computed for each L-R pair documented in the LR*db* database. We typically use *k*_1_ = 5 because each random dataset yields a large number of random correlations, one for each L-R interaction in LR*db* with ligands and receptors in the matrix *A*. We pool all correlations to estimate the null distribution. For some datasets, a censored normal distribution (correlations lie in [−1; 1]) provides an accurate fit (Figure S1A). With 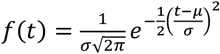 the density function of a *N*(*μ,σ*) distribution, the density of the censored normal distribution is 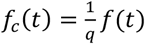, with 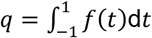 The parameters *μ* and *σ* are estimated with a maximum-likelihood (ML) approach. However, in some cases, it is necessary to use a mixture of two censored normal due to a slight asymmetry (Figure S1B). A mixture of two normal distribution density functions is given by 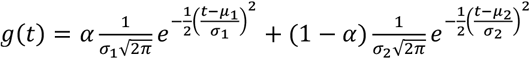, and the censored mixture distribution is obtained as above: 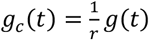, with 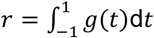. The five parameters *α*,*μ*_1_,*σ*_1_,*μ*_2_,*σ*_2_ are estimated by ML. We also implemented a purely empirical distribution and a Gaussian kernel-based empirical distribution (Figure S1C).

In ST, random correlations tend to be more asymmetric (biased towards positive values) and heavy-tailed. Besides the previously described empirical models, we found that censored stable distributions fitted such data accurately (Figure S1D). Stable distributions are a family of distributions that include normal, but also heavy-tailed distributions, such as Cauchy distributions. The stable distribution density function is given by (Nolan representation) 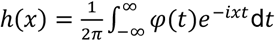, with *φ*(*t;α,γ,δ*) = *e*^*itsδ*-|*γt*|^*α*^(1-*iβ*sgn(*t*)φ)^, where sgn(*t*) is the sign function and 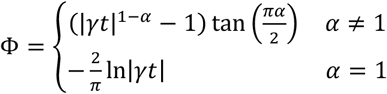. The censored stable density is obtained with 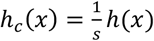 and 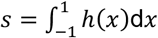. The four parameters *α,β,c,μ* are estimated by ML. The stable density and cumulative distribution functions are provided by the R package stabledist.

The R-T null distribution is obtained in a similar manner. For each randomized dataset, we consider all combinations of receptors (from the *LRdb* database) and downstream pathways. For each combination, we identify the target genes and we compute their Spearman correlation with the receptor. We then pool all correlations for all receptors and pathways. We repeat this for *k*_2_ randomized datasets, again pooling all correlations. We typically use *k*_2_ = 2 because each iteration yields >100,000 random correlations. In the special case of the censored stable distribution, subsampling is used to avoid endless computations by randomly selecting the same number of random correlations as obtained for the L-R null distributions. As the L-R null distribution and the R-T distribution are usually very similar, BulkSignalR offers the possibility to use the L-R null distributions for R-T to save training time. We do not recommend this option for accurate computations, but it may be convenient for preliminary analyses, especially when using stable distributions where parameter estimation can take up to 15 minutes on a powerful processor. We tried the expectation-maximization algorithm implemented in the R package alphastable, but it did not fit our distributions and required comparable computing time (data not shown).

The BulkSignalR parameter training algorithm automatically chooses a model among the censored normal, censored mixture of two normal, and Gaussian kernel-based empirical models.

However, the user can impose a model manually, including purely empirical and censored stable models. A control plot displaying the empirical histograms of (L-R) and (R-T) correlations and the chosen fitted model can be generated by BulkSignalR training function (Figure S1). The model selection algorithm is:

- Compute *χ*^2^ statistics between a random correlation histogram with equally sized bins (bin width = 0.05) and the censored normal, censored mixture of normal, and Gaussian kernel-based empirical models.
- Select the censored Gaussian if its *χ*^2^ is not worse than 1.25 times the censored mixture *χ*^2^ and 2 times the Gaussian kernel *χ*^2^.
- Select the censored mixture if its *χ*^2^ is not worse than 2 times the Gaussian kernel *χ*^2^
- Select the Gaussian kernel-based empirical model, otherwise.

### Statistical model

Under the null hypothesis, we assume that L-R and R-T correlations are independently and identically distributed after their respective null distributions. Accordingly, we estimate independently and multiply the significance of the L-R correlation and the set of R-T correlations. We obtain the L-R correlation significance directly from its null distribution. To allow the search of antagonist ligands (see Results), L-R significance computation depends on its sign. If *F*(*r*) is the cumulative distribution function (CDF) of random L-R correlations *r*, then we use P-value=1 - *F*(*r*) for *r* ≥ 0, and P-value=*F*(*r*) otherwise.

It is very difficult to assess the activity of the pathways downstream of the receptor in full generality, considering pathways of all possible sizes and topologies, and also RNA-seq, DNA chip, or proteomic data. Moreover, we wanted BulkSignalR to learn from the expression dataset directly, without any manual intervention, and to be applicable to datasets of virtually any size. Therefore, we opted for a simple but robust approach. Once the target genes *g_1_*,…,*g_N_* of a pathway *pw* downstream a receptor *R* are identified, we can compute the Spearman correlations between *R* and each of *g_i_*. We use order statistics (making the assumption of independence between R-T correlations under the null hypothesis) to integrate the information provided by several target genes, and also to take into account branches of a large pathway that are not all active, or regulatory mechanisms that cannot be detected with the used technology, *e.g*., posttranslational modifications in transcriptomic data. Namely, all the R-T correlations are sorted by increasing order and the *k*^th^ percentile is used. With *k* = 100 we only check the best correlation and with *k* = 75 we check the 75^th^ percentile of all correlations. By construction, the order statistics integrates the pathway size and the correlation strengths. For a *k*^th^ percentile correlation *r_k_*, the probability to observe a Spearman correlation *R* ≤ *r_k_* is given by the CDF of the chosen censored distribution *F_R_*(*r_k_*). The order statistics CDF is given by a binomial distribution 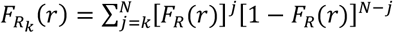 and P-values are obtained from *F_Rk_*(*r*). Multiple hypothesis correction is applied using the R package multtest with the Bejamini-Hochberg method as default method.

### Gene signature scoring

We implemented a simple scheme where each gene is normalized separately across samples to obtain z-scores. The score of a gene signature is the average of the z-scores of all included genes.

### LRI association with cell types

We assume that two matrices of gene signature scores are available, a *t*×*m* matrix *C* for cell types and a *r*×*m* matrix *I* for LRI activity. That is, *t* different cell type signatures and *r* distinct LRI signatures were scored over the *m* sample of the dataset. To associate the LRI on row *k* of the matrix *I* with one or several cell types, a preliminary filter requires a minimum Spearman correlation between a cell type signature and the LRI signature *I_k__i_* (default = 0.25). If any cell type passes this filter, the selected cell types are used to construct a regularized linear model with the LASSO and by imposing non-negative coefficients (otherwise default parameters of the glmnet R library were used to optimize the weight *λ* of the penalty term). If we denote *P_t_* the set of cell types with non-zero weights and *P_t_* ≠ ø, the linear model ∑_*j∈p*__*t*_^*α_j_C_j_*^. approximates *I_k_r__*. We remove all cell types with a low weight *α*_7_ in the model (< 0.1 by default) to obtain a new set of cell type indices 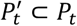. The model is considered valid provided its correlation with LRI activity, *i.e*., Spearman correlation between 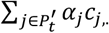 and *I_k_*, is sufficient (> 0.35 by default).

### Association with tissue areas

Users can choose among a default statistical non-parametric model (Kruskal-Wallis for global association and Wilcoxon for each area *versus* all the others), a parametric Gaussian model (ANOVA and t-test), Spearman correlations, and the coefficients of determination (r2) of linear regressions. In the two last models, tissue regions are represented by their characteristic functions (1 = inside the region, 0 = outside).

### Implementation

BulkSignalR is implemented in R following an S4 object-oriented approach. The package follows the Bioconductor standards (submission pending) and is available from GitHub.

### Datasets and their preparation

All the examples provided in this article were based on public data made available by their respective authors, but for the CRC-LM spatial data that were generated by us (see below). We downloaded TCGA RNA-seq data (gene read counts) from the BROAD Institute GDAC at firebrowse.org. The frontal cortex transcriptome data were from the Genotype-Tissue Expression (GTEx) Project v8 (2017-06-05) RNASeQCv1.1.9_gene_reads_gct. Lung cancer cell line transcriptome (RNA-seq) data were from DepMap (15) 22Q2 Portal.

BulkSignalR includes a data preparation function that eliminates non- or barely expressed genes/proteins and performs common normalization procedures. We chose default parameters for RNA-seq data: genes were retained if a minimum read count of 10 was found in at least 10% of samples (percentage and minimum value can be changed to adapt to different data). The default normalization is upper quartile, but total count is available as well. Pre-normalized data can be used to allow filtering and normalization according to more advanced strategies. We processed datasets with default parameters unless otherwise specified. For the datasets presented in Supplementary Information, refer to Supplementary Methods. For the DepMap lung cancer cell line data, we imposed a minimum read count of 1 and log-transformation.

### Pseudo-receiver operating characteristics (ROC) curves

To estimate true and false positives (TPs and FPs), we applied BulkSignalR to the original data matrix *A* and also randomized matrices (see above for the randomization procedure). By varying a threshold on the Q-values we obtained estimates of the number of FPs from the randomized data for that specific Q-value. The corresponding number of TPs was estimated by the number of selected LRIs at the same Q-value from the original matrix *A* minus the number of FPs. We named these curves pseudo-ROC curves because the estimates are obtained in the absence of an exact reference. To estimate their variability, we generated 100 randomized expression matrices *A* per dataset. We also generated pseudo-ROC curves for single-cell scores (similar to what done by ICELLNET and CellPhoneDB). We computed ICELLNET-like scores using normalized data according to the ICELLNET original publication (11). We obtained CellPhoneDB-like P-values from 1,000 shuffled datasets. We did not implement any specific treatment for multimeric receptors, thus departing from the original CellPhoneDB and ICELLNET implementations. We estimated the pseudo-ROC curve variability based on 50 randomized expression matrices *A* in this case.

### Comparison with other ST software tools

We used BulkSignalR with default parameters except min.count = 1, method = “TC”, min.prop = 0.01 when calling the method prepareDataset(), min.positive = 2 when calling the method learnParameters(), and min.cor = −1 when calling the method initialInference().

For stLearn, we followed the steps provided by the Cell-Cell interaction tutorial in the stLearn documentation website. We used the function st.tl.cci.run() with distance = 0 to compute LRIs in the within-spot mode, and we set n_pairs (the number of random pairs generated to compute the background distribution) to 1,000. We did not apply any P-value correction for multiple hypotheses. We replaced the stLearn L-R database by LR*db* (Cabello-Aguillar 2020 NAR). Then, we exported all L-R and the associated P-values from data.uns[‘lr_summary’] and data.obsm[‘p_vals].

We used CellPhoneDB v3 to compute inferences of LRIs according to the documentation provided by the CellPhoneDB website. CellPhoneDB database v4 was used. We set the method parameters to “statistical analysis”. We incorporated spatial information of cell types via the microenvironments file.

We used SpaTalk to directly infer cell-cell communications, thus skipping the deconvolution mode as described in its Wiki documentation. We called the function createSpaTalk() with the following parameters: if_st_is_sc = FALSE, spot_max_cell = 1. We defined the major cell type at each spot according to the data released by the authors of the different datasets. We used the LR*db* database as the core database for LRI, thus replacing the native database. We called LRIs with downstream targets using the function find_lr_path().

### Liver metastasis patient material

The ICM Translational study committee, Montpellier, approved the present study. In accordance with the French law, patients who did not oppose to the use of their material for research purposes provided consent (opting-out rule). Four CRC-LM samples were used in the present study.

### Liver metastasis spatial gene expression analysis

First, we analyzed the quality of RNA extracted from selected formalin-fixed paraffin-embedded (FFPE) CRC-LM samples by evaluating the percentage of fragments with length > 200 nucleotides (DV200). Briefly, we cut one-two 10 μm-thin FFPE tissue sections, followed by dewaxing and lysis. We extracted and purified RNA using the High Pure FFPE RNA Isolation Kit (cat. N. 06650775001, Roche), and analyzed the samples by microfluidics-based automated electrophoresis (Bioanalyzer; Agilent). All tested samples had a DV200 > 50%. Then, we used serial sections from the same tissue blocks, stained with Hematoxylin Eosin Saffron (HES), to identify an area of ~6 x 6 mm containing liver metastasis, in order to provide optimal coverage of the 6.5 x 6.5 mm capture area on the Visium slide. Following macro-dissection, we fixed 10 μm-thin sections of the selected areas on the Visium slide, followed by dewaxing, HES staining, imaging, and decrosslinking according to the manufacturer’s instructions (10X Genomics Visium Spatial Gene Expression for FFPE kit, Human Transcriptome, Protocol n. GC000408 and GC000409). Then, we prepared Visium libraries according to manufacturer’s instructions (Protocol n. GC000407). After sequencing using the NovaSeq 6000 system (S1 flowcell; Illumina) to obtain ~50,000 reads per spot, we used the Space Ranger Software Suite (version 1.3) for sample demultiplexing, alignment to the human probes, barcode assignment of each spot, and gene counting by unique molecular identifier counts. Data will be available at GEO upon publication.

### Multiplexed immunofluorescence

After deparaffinization in xylene, rehydration in a series of methanol dilutions (see above), and antigen retrieval in AR6 buffer (Perkin Elmer, Waltham, MA, USA; cat. n. AR600250) in a pressure cooker for 10 min, we incubated FFPE CRC-LM tissue sections in serum-free blocking solution (Agilent-Dako, Santa Clara, CA, USA; cat. n. X0909) at room temperature for 30 min followed by incubation with the anti-epidermal growth factor receptor (EGFR) primary antibody (cat. n. 4267, Cell Signaling) at 4°C overnight. Then, we washed slides in TBS−0.01% Tween 20 and incubated them with Histofine MAX PO Multi (Nichirei, Tokyo, Japan; cat. n. 414152F) secondary antibody at room temperature for 30 min. We performed staining with the TSA Coumarin system (Akoya, cat. n. NEL703001KT). Next, after antibody stripping with the AR6 buffer in a pressure cooker for 5 min, and antigen blocking for 30 min, we incubated sections with three antibody sets (each set containing three antibodies), by repeating the staining, stripping and blocking steps for each antibody. Set 1 (structure) included: anti-CD31 (IR61061-2, Dako), anti-pan-CK (GA05361-2, Dako) and anti-SMA (GA61161-2, Dako) antibodies. Set 2 (decorin) included: anti-phosphorylated p44/42 MAPK (Thr202/Tyr204) (4376, Cell Signaling), anti-cMET (8198, Cell Signaling), and anti-DCN (AF-143, R&D Systems) antibodies. Set 3 (cadherin) included: anti-phosphorylated ERK1/2, anti-cMET (same antibody as in set 2), and anti-CADH1 (GA05961-2, Dako) antibodies. The secondary antibody was the same for most antibodies (Histofine MAX PO Multi), but for the anti-DCN antibody that was detected using Histofine MAX PO G (414162F, Nichirei Bio). For staining we used the Opal system (Perkin Elmer, cat. n. NEL810001KT). After mounting using VECTASHIELD^®^ Vibrance Mounting Medium without DAPI (Vector, Burlingame, USA), we visualized staining using a Thunder microscope (Leica, Wetzlar, Germany).

## RESULTS

### BulkSignalR approach and design

Most single-cell tools infer LRIs by relying only on the ligand *L* and the receptor *R* abundance because they have access to separate data for each cell population. With bulk data, the observed abundance of transcripts (or proteins) is the net contribution of different cell types, each of which represents an unknown proportion of the analyzed tissue. Therefore, we hypothesized that LRI inference using bulk data should rely on additional modeling steps. Specifically, we modeled the consequence of a putative LRI by considering the participation of *R* in biological pathways and the regulation of the genes targeted by these pathways. Furthermore, as we did not assume any knowledge about the different samples, such as clusters harboring similar profiles, we modeled relationships between ligands, receptors, and downstream pathways based on Spearman correlations across the whole dataset. Consequently, BulkSignalR searches for triples (*L,R,pw*) where *pw* is a pathway downstream of *R* (Figure 1A). *L, R*, and *pw* must display correlated activities to be deemed significant by our statistical model. Potential LRIs are taken from the LR*db* database (7), while pathways originate from Reactome (16) or GO biological processes (17) regarded as pathways. To identify genes reporting on a pathway activity, which we called target genes in general, we exploited the topology of a reference network that includes Reactome and KEGG (18) interactions. Target genes include the targets of transcription factors of the pathway. By default, when *R* is part of a complex, we add the other complex members to the list of targets because their expression should be correlated with *R* expression to maintain the complex stoichiometry. BulkSignalR uses a statistical model to assess the significance of all possible triples (*L,R,pw*) based on the null distributions of L-R and also R-T correlations, pathway sizes, and total number of target genes (see Materials and Methods). Importantly, BulkSignalR computes null distributions of correlations from the (randomized) dataset directly, and the statistical model combines all correlation values independently from the sample number. This avoids the issue of very small, but highly significant correlations in large datasets and allows the analysis of small cohorts.

**Figure 1.**
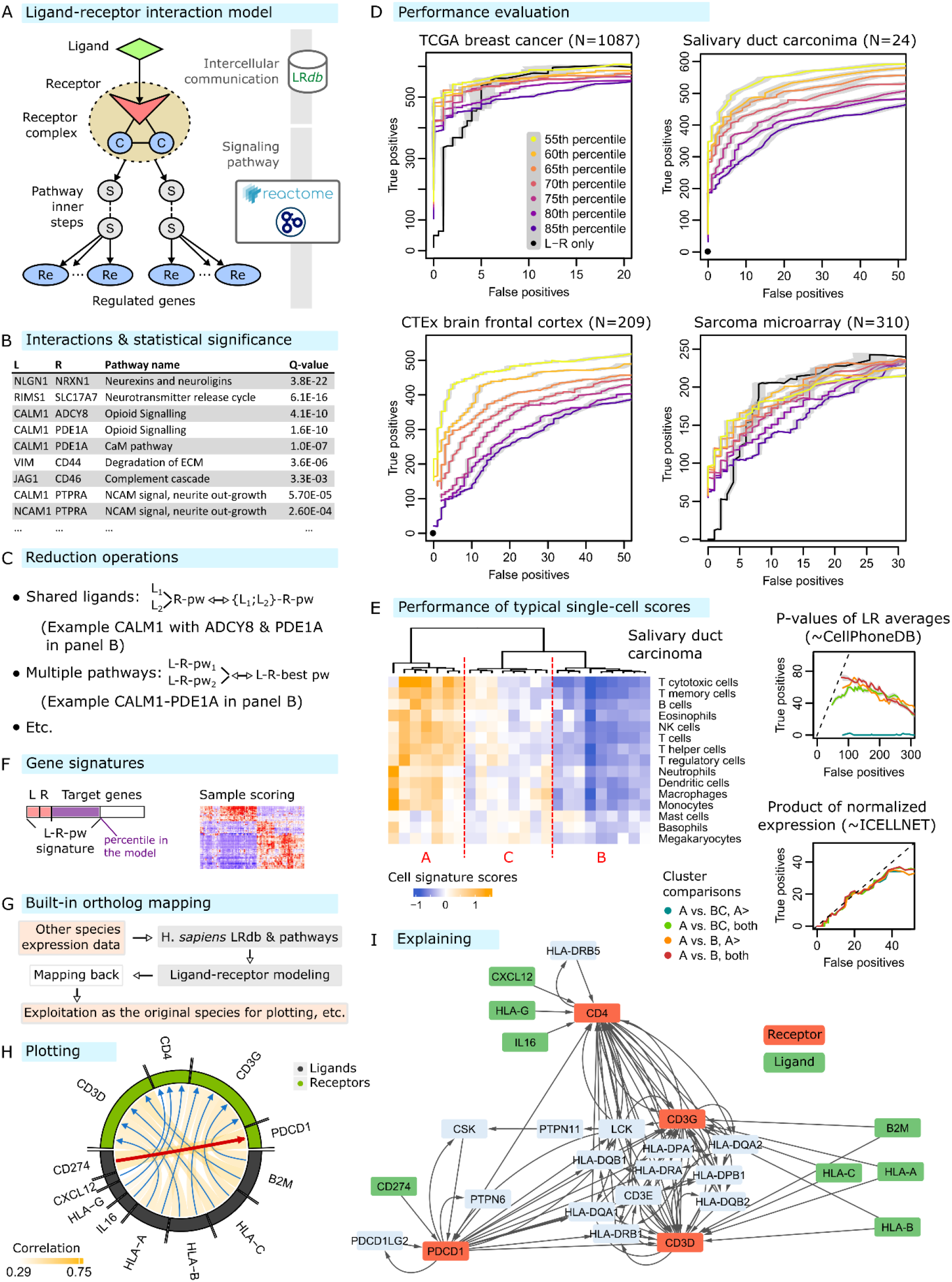
BulkSignalR overview. (**A**) Model of a LRI with a pathway downstream the receptor. The pathway activity in terms of correlation with the receptor is reported by target genes (in blue) that include the other members of the receptor complex (denoted by C) and regulated genes (denoted by Re). (**B**) Example of BulkSignalR output. (**C**) Examples of reduction operations at the receptor level and at the best pathway level. Other reduction operations are available. (**D**) Representative pseudo-ROC curves showing the effect of using the pathway information at various depths (depth increases with smaller percentiles (0.55 = 55^th^ percentile). RNA-seq transcriptome data of 1087 TCGA breast invasive carcinoma samples (BRCA), 24 salivary duct carcinoma (SDC) samples (6), and 209 brain frontal cortex samples retrieved from GTEx v8; microarray transcriptome data (Affymetrix) of 310 sarcoma samples (48). In the SDC and GTEx brain cortex datasets, no individual L-R correlation alone did not even reach 5% of significance and a simple dot at (0,0) was drawn to indicate this. (**E**) Performance of simple single-cell-inspired scores when samples display clear differences in cellular composition. We considered clusters A *vs*. B∪C and A *vs*. B to look for increased scores compared with the randomized expression data in A or in both directions. (**F**) Construction of a gene signature for a L-R-pathway triple, and computation of the weighted average of z-scores. Multiple L-R-pathway signature scores can be compared among samples (heatmap). (**G**) In BulkSignalR, the reference database is of human origin, but an integrated ortholog mapping tool allows using it for virtually any species. (**H**) Example of graphical display of LRIs limited to a chosen pathway (PD-1 signaling in the SDC dataset). (**I**) Representation of the LRIs of the chosen pathway (PD-1 signaling in the SDC dataset) with the shortest paths to target genes.

The results of BulkSignalR statistical analysis are summarized in a table that contains putative LRIs, pathways including the receptors, and statistical parameters (Figures 1B and S2). Due to redundancies in pathway definitions that often occur at different levels of detail, the same interaction can appear in multiple pathways. Moreover, some ligands can have more than one receptor and *vice versa*. Therefore, we introduced different reduction operations to limit redundancy, or to emphasize the ligand, the receptor, or the pathway (Figures 1C and S3). Such operations can be chained. Reduction to the pathway that yields the best P-value for each LRI results in a table with unique interactions. Using this reduction, we estimated the TP and FP rates in order to obtain pseudo-ROC curves (Figure 1D). Such curves allowed varying the depth at which target gene correlations are assessed in different datasets, which revealed that it was advantageous to exploit the pathway downstream the receptor, and that the L-R correlation on its own did not provide a useful inference mechanism. Indeed, the performance of sole L-R correlation was limited or did not detect any significant LRI. Although we could evaluate the pathway activity by considering all its target genes, large pathways may comprise multiple branches, and target genes may be up- or down-regulated. As a compromise, we did not go deeper than the 45% of the highest correlation (55^th^ percentile in Figure 1D).

The number of LRIs found by BulkSignalR was lower compared to what was obtained when comparable single-cell datasets are available. Nonetheless, as expected, only the interactions found using bulk data were enriched in low-abundance molecules (Figure S4).

As explained in the Introduction, single-cell scores have been proposed to infer LRIs from bulk data, although they require separate datasets for the ligand-secreting cells and the receptor expressing cells. We investigated whether such scores could be adapted when no upfront cell separation is available. To this aim, we selected a salivary duct carcinoma (SDC) dataset (6) that included well-separated clusters of samples with limited intra-cluster variability. The scoring of common immune cell gene signatures (Table S1) revealed such clusters (Figure 1E). We then compared the clusters to identify LRIs that were enriched in one cluster as a selection mechanism. At the time of writing, the LR*db* database included 3,249 putative LRIs. Due to the necessary presence of both ligand and receptor in the dataset, and the default requirements of BulkSignalR analysis (L-R correlation > 0.25, minimum four target genes found in pathways of sizes between 5 and 200), we could evaluate 777 LRIs with BulkSignalR. We submitted those 777 LRIs to adapted single-cell scores. We used a score based on ICELLNET (a product of average values) with specific data normalization (11), and another one based on CellPhoneDB (P-values of the ligand and receptor average expression by sample random permutations across clusters). We compared cluster A *vs*. BuC, respectively A *vs*. B, to obtain strong, respectively extreme, differences in tumor immune infiltrate abundance (Figure E). Furthermore, we selected scores that were higher in the cluster with most immune cells or in one of the two compared clusters. This resulted in four pseudo-ROC curves for each score (Figure 1E), none of which performed better than random selection.

To compare samples and search for differential activity of LRIs is an important functionality. Therefore, we introduced the notion of gene signature to reflect the overall activity of the interactions, including downstream pathway targets (Figure 1F). The expression values of different genes (or proteins) are transformed into z-scores and a weighted sum defines the score. By default, the average ligand and receptor z-scores accounts for one half, while the target genes included in the statistical model (correlations above chosen percentile) account for the other half of the score, with equal individual weights. Gene signatures and scores remain compatible with any combination of reduction operations due to BulkSignalR software design. To facilitate the analysis of non-human datasets, we integrated a generic ortholog mapping mechanism that allows users to virtually work with any species (Figure 1G). We designed BulkSignalR with the aim of proposing a user-friendly tool. Therefore, only few lines of code, accessible to basic R users, are sufficient to perform a complete analysis and generate informative outputs (Figure S5) including graphical representations (Figures 1HI and S6).

### Autocrine communications in lung cancer cells

For the first application of BulkSignalR, we analyzed bulk transcriptomic data from 206 lung cancer cell lines from DepMap (15). We reduced BulkSignalR output to the best pathway for each LRI to obtain only unique interactions, and we imposed a FDR < 0.1% (full output in Table S1). We obtained and scored gene signatures (Figure 2A). Cell lines originating from the two main histologic subgroups, small-cell lung cancer (SCLC) and NSCLC, harbored distinct autocrine communication patterns. Mesothelioma-derived cell lines were close to those of NSCLC-derived cell lines, although they constitute a different entity. This similarity may simply reflect the absence of microenvironment in cultured cell lines. The presence of only two lung carcinoid cell lines in the dataset prevented drawing any conclusion on this subgroup.

**Figure 2.**
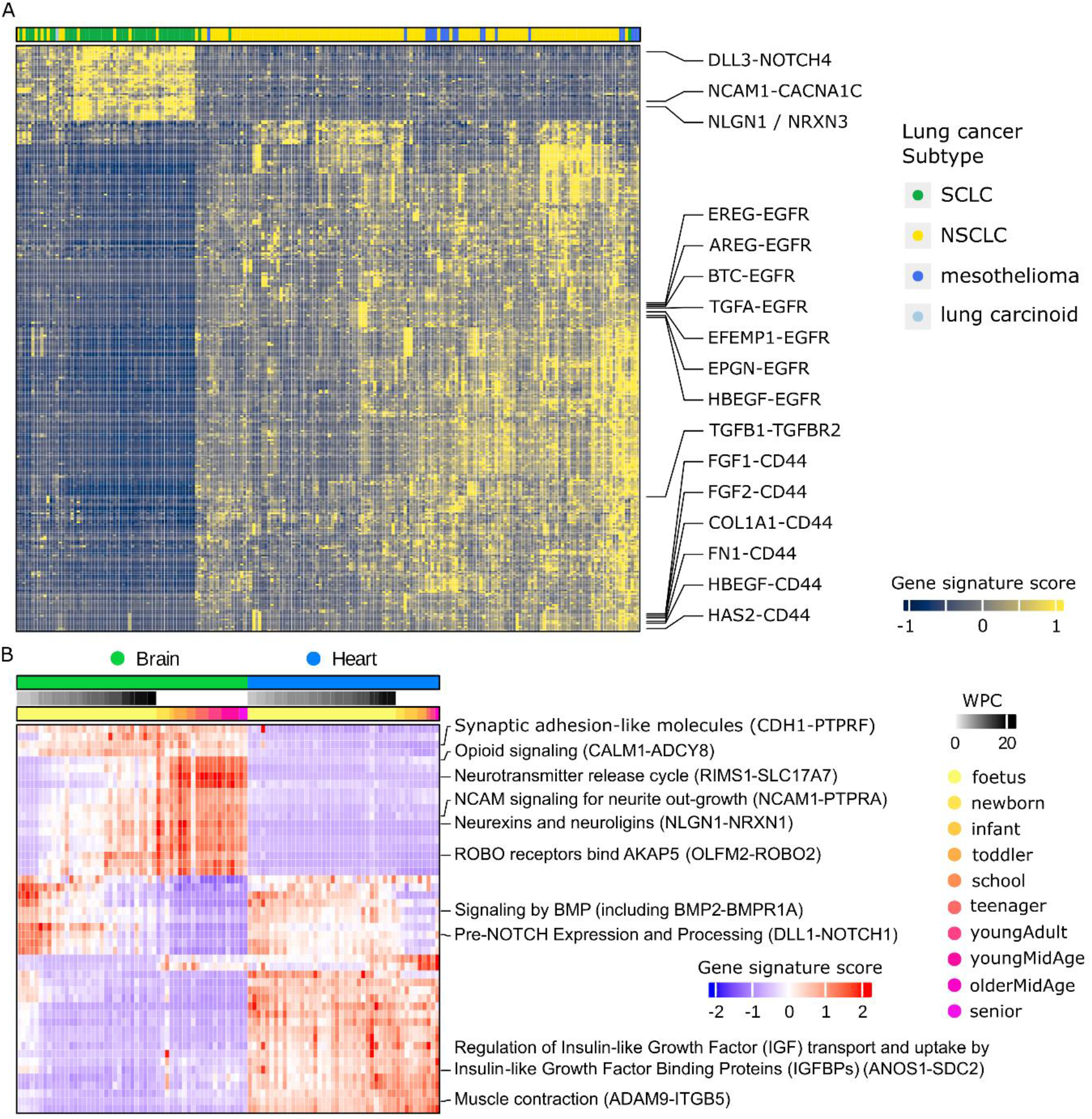
Bulk data analysis with BulkSignalR. (**A**) Autocrine communications in the DepMap lung cancer cell line dataset. Analysis was performed at the LRI level. Representative interactions are indicated (FDR < 0.1%). (**B**) Analysis (pathway level) of transcriptome data of human brain and heart samples at different development and adult life stages (32). Representative pathways are indicated with one LRI example in brackets. Pathways were taken from Reactome. FDR < 0.1%. WPC = weeks post-conception.

In the NSCLC cell lines, we obtained the highest scores for LRIs that involved EGFR. This is in line with its increased activity in more than half of patients with NSCLC (19). We identified specific interactions between EGFR and its seven activating ligands (20) (Figure 2A). It is important to be able to identify each interaction because different ligands may be associated with functional differences (21). The pathways associated with these interactions included EGFR downregulation, clathrin-mediated endocytosis, general ERBB2-related signaling processes, signaling by PTK6, and signaling by non-receptor tyrosine kinase (Table S2). An example of ligand-specific signaling was related to the heparin binding EGF-like (HBEGF)-EGFR interaction, for which we identified additional pathways, *e.g*., NOTCH3 Activation and Transmission of Signal to the Nucleus (Table S1). Previous studies already reported the crosstalk between EGFR and the Notch pathway in NSCLC (22), and the important specific role of the HBEGF ligand (23). LRIs harboring a prognostic value also were enriched in NSCLC cell lines. For example, TGFB1-TGFBR2 is involved in TGF-β receptor signaling associated with the Epithelial-Mesenchymal Transition (EMT) pathway (24, 25). Other LRIs involved the CD44 receptor, which is often described as a cancer stem cell marker. In NSCLC, CD44 and its isoforms have been associated with poor prognosis and tumor invasion (26).

In agreement with the fact that SCLCs are high grade neuroendocrine tumors, we found several LRIs related to neurexins and neuroligins, for instance NXPH1-NRXN1 and NLGN1-NRXN3. Interestingly, the NRXN1 receptor is considered a novel potential target of antibody-drug conjugates against SCLC (27). The NCAM1 ligand, which is a surface marker for SCLC (28), interacted with the PTRA and CACNA1C receptors. These interactions are needed to activate signaling for neurite outgrowth, thus contributing to the neuroendocrine phenotype. Lastly, other LRIs suggested the potential activation of Notch-related signaling, for instance the DLL3-NOTCH4 interaction. In SCLC, Notch signaling may have tumor suppressive or promoting activity and is a candidate biomarker of the response to immune checkpoint blockade (29). Recent studies investigated treatments targeting DLL3 in recurrent SCLC (30), and in high-grade pulmonary neuroendocrine tumor-initiating cells (31).

### Summarizing at the pathway level

In the analysis of lung cancer bulk data, we found LRIs that implicated a single receptor or related to a single pathway. BulkSignalR capacity to reduce data can be exploited to investigate a dataset at the pathway level. We illustrate this procedure using transcriptomic data from human brain and heart during development and adult life (32). We started by reducing the output at the pathway level. To this aim, we pooled together all receptors implicated in a given pathway, followed by pooling of all their ligands. In this way, that pathway was associated with a meta-ligand and a meta-receptor instead of many single interactors (Figure S3): a standard triple (*L,R,pw*) becomes ({*L*_1_,…,*L_N_*},{*R*_1_,…,*R*_M_},*pw*). If two different pathways were associated with the same meta-ligand and-receptor, we performed a reduction to maintain only the best pathway chaining BulkSignalR chained reduction operations (Figure S6).

The pathway-level analysis (Figure 2B) identified tissue-specific, and also shared pathways and LRIs, including some that are development stage-dependent. Brain-specific pathways included Synaptic adhesion-like molecules (including the LRIs CDH1-PTPRF and PTN-PTPRS), and at later stages Opioid signaling (including CALM1-ADCY8 and CALM2-PDE1A) and Neurotransmitter release cycle (RIMS1-SLC17A7). NCAM signaling for neurite outgrowth (NCAM1-PTPRA and PSPN-GFRA2), Neurexins and neuroligins (NLGN1-NRXN1 and NXPH1-NRXN1), and ROBO receptors bind AKAP5 (OLFM2-ROBO2) were additional brain pathways that start *in utero* and show maximal activity after birth. Shared developmental pathways were Signaling by BMP (BMP2-BMPR1A and GDF11-ACVR2B) and Pre-NOTCH Expression and Processing (DLL1-NOTCH1, DLL4-NOTCH4, and JAG1-NOTCH1). Regulation of insulin-like growth factor transport and uptake by insulin-like growth factor binding proteins (ANOS1-SDC2 and FGF2-SDC2) and Muscle contraction (ADAM9-ITGB5, ANXA1-DYSF, and COL1A1-ITGA1) are examples of heart pathways. The complete list of all LRIs and pathways is provided in Table S1.

### Relating ligand-receptor interactions to cell types

In the single-cell paradigm, LRI inference comes with the knowledge of which cell types express the receptor and the ligand. As such information is not directly accessible in bulk datasets, we implemented an algorithm to predict cell type-LRI associations. This requires scoring a set of cell type gene signatures in all samples. It can be achieved with a simple z-score average-based function provided by BulkSignalR, or with more advanced cellular deconvolution tools the output of which can then be imported in BulkSignalR. By comparing cell type and LRI gene signature scores (Figure 3A), we build a sparse linear model in which LRI activity is linked to cell type abundances by the following equation: LRI *k* activity 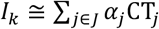 for a small set *J* of cell types CT_*j*_. Due to the intrinsic limitations of bulk data and the absence of complete LRI reference, BulkSignalR cannot predict whether a given cell type expresses the ligand, the receptor, or both. Using the SDC dataset, we scored cell type signatures for common tumor-infiltrating cell types: immune cells, endothelial cells, and fibroblasts (Table S1). We summarized cell communications by summing the weights of the cell types in all LRI activity models as above, *i.e*., by summing the *α_j_* values (Figure 3B). We noted stronger communications within the stromal (endothelial cells and fibroblasts) and immune components of the tumor, as expected. By selecting all LRIs that were exclusively associated with stromal cells or with immune cells, we found that they were involved in very relevant pathways (Figure 3B). Immune pathways included well-known immune checkpoints, in agreement with the strong immunosuppressive microenvironment of immune cell-infiltrated SDCs. We experimentally identified these immune checkpoints and some of the implicated cell populations in a previous study on the SDC landscape (6), and found a good agreement between our previous results and the BulkSignalR associations (Figure 3C). In the absence of available ground truth, we relied on synthetic data to obtain a more general performance estimation of cell type assignment. By randomly picking 1, 2, or 3 cell line signature scores and adding Gaussian noise, we generated correct cases, *i.e*., artificial LRI signature scores that should be assigned to the randomly picked cell lines. The noise standard deviation *σ* in real data varies between 0.2 and 0.4. We computed the TP rates (TPR) and true negative rates (TNR) using the small SDC and the large TCGA BRCA cohorts (Figure 3D). We also generated randomized data that should not be assigned to any cell type by picking LRI signature scores and shuffling 10, 25, or 50% of the values (Figure 3D). The obtained TNR and TPR showed that the algorithm avoided wrong assignments, but its sensitivity to detect complex relationships with three cell types decreased with increasing noise, especially in the smaller dataset.

**Figure 3.**
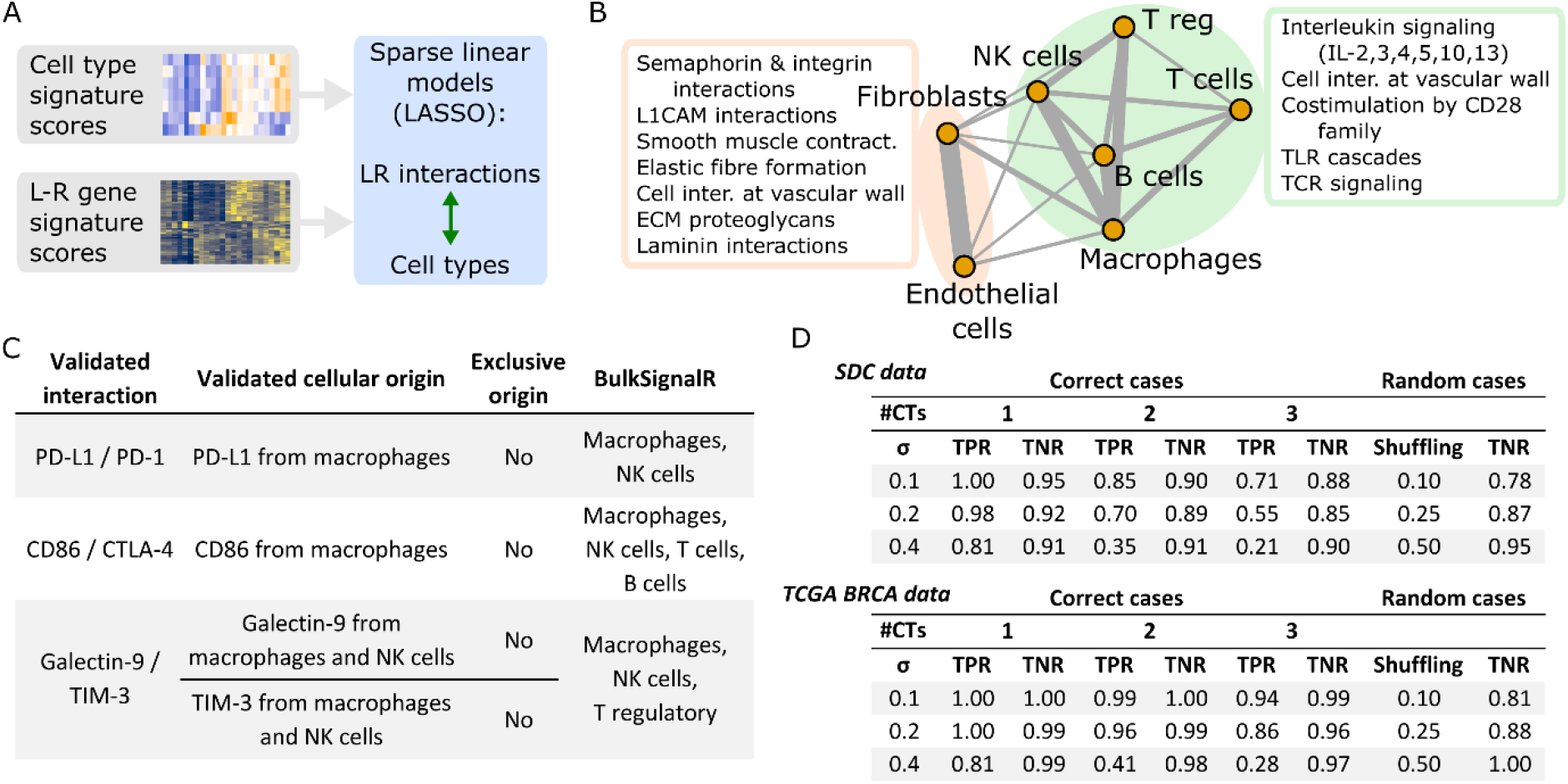
Assigning cell types to interactions. (**A**) Using the gene signature scores for cell types and the gene signature scores for LRIs, mathematical models are built to predict the L-R scores from a minimum set of cell type scores. (**B**) For the SDC dataset, we represented the total weights of each cell type contribution to LRIs (network). Left: most recurrent pathways related to LRIs only between fibroblasts and endothelial cells. Right: most recurrent pathways for interactions only between immune cell types. These pathways include important immune checkpoint molecule interactions: (PD-L1)-(PD-1), (Galectin-9)-(TIM-3), and (CD80/86)-(CTLA-4). (**C**) Immunofluorescence-validated interactions and cell types in SDC (6) and BulkSignalR predictions. (**D**) True positive rate (TPR) and true negative rate (TNR) of the cell type association algorithm for synthetic data. Correct cases include interactions involving 1, 2, or 3 cell types and increasing Gaussian noise N(0,*σ*). Random cases were obtained by shuffling 10, 25, or 50% of data. Synthetic models were generated using a small (SDC) and a large (TCGA breast cancer, BRCA) dataset. In total, 100 synthetic models were constructed for each configuration (correct/random, σ, shuffling rate, #CTs = number of cell types).

### Other functionalities and applications

Our model of LRI (Figure 1A) implies significant positive correlations between the receptor and a downstream pathway as well as correlations between ligand and receptor. By default, BulkSignalR does not consider L-R correlations < 0.25. However, it is possible to impose a different minimum, particularly −1, and to find a number of (*L,R,pw*) triples with strong P-values, but negative L-R correlations. This case is properly handled by our statistical model and suggests that such ligands may have an inhibitory action. We reanalyzed the SDC dataset by imposing FDR < 0.1% and L-R correlation > 0.25 in absolute value. We identified 424 unique LRIs, 361 positive and 63 negative (Table S1). By focusing on the Notch pathway, which is deregulated in many tumors, we found common activators (DLL1, DLL4, DLK1, DLK2, JAG1, and JAG2) with strong P-values and positive correlations with one or several of the four Notch receptors. We also found several interactors that were negatively correlated with Notch receptors, for instance MFNG (correlation −0.52 with NOTCH1 and NOTCH2), a glycosyltransferase that modulates Notch activity by modifying O-fucose residues at specific EGF-like domains (33). DLL3, which can inhibit Notch (34), also displayed negative correlations (−0.26 with NOTCH3 and, just below our 0.25 threshold, −0.245 with NOTCH4) as well as PSEN1 (−0.50 with NOTCH1, −0.27 with NOTCH2, and −0.41 with NOTCH3) and UBA52 (−0.51 with NOTCH1). It is worth noting that searching for negatively correlated L-R pairs is much more prone to FP (Figure S7) because L-R databases are dominated by activating LRIs. Therefore, this procedure should be considered as exploratory and the output needs to be experimentally validated.

NicheNet (35), a single-cell tool, exploits an integrated molecular interaction network that includes LRIs to relate user-chosen gene sets to ligands that might drive their expression according to the network. The authors tested this functionality using the 100-gene signature proposed by Puram (5) for a partial EMT transcriptional program taking place at the invasive front of head and neck squamous cell carcinoma (HNSC). As NicheNet integrated reference network and our method to link ligands to receptor-pathway target genes are conceptually similar, we use BulkSignalR inferences to provide a similar functionality, but for bulk data. Figure S8 describes its application to investigate the partial EMT program using the TCGA HNSC bulk transcriptome dataset (*n* = 500).

In Supplementary Information, we describe the analysis of DepMap transcriptomic and proteomic data of breast cancer cell lines (Figures S9-S12), and we use a mouse dataset to illustrate the built-in ortholog mapping functionality (Figures S13-S15). Moreover, BulkSignalR allows replacing LR*db*, its native L-R database, with a user-provided equivalent or adding user-chosen L-R interactions.

### Investigating spatial transcriptomics data with BulkSignalR

We then applied BulkSignalR to several multicellular resolution datasets (Visium). BulkSignalR functions were used unchanged, we only set some parameters to adapt them to the much reduced data dynamics and large number of spots (Figure S16). We started with a triple-negative breast cancer (TNBC) dataset (36) the overall structure of which is presented in Figure 4A. We identified 224 LRIs (Table S1 and Figure S17), some of which we briefly present here to show that BulkSignalR identified relevant interactions. Most LRIs were associated with the stroma or with the invasive cancer tissue. In the stroma, we found LRIs that implicated adhesion molecules, *e.g*., CAM1, VCAM1, and several integrins, *i.e*., ITGB1, ITGB2, ITGB7, ITGAL, ITGAX, as well as immune-related L-R complexes, *e.g*., B2M-CD3D, B2M-(HLA-F), (HLA-A)-CD3D, and IL16-CD4. The presence of activated interactions related to antigen presentation (B2M; HLA-A/B) was consistent with findings by the authors of this dataset (36). Figure 4B shows the spatial distribution of the LRI between the metalloproteinase MMP9 and the integrin ITGB2. MMPs are important TME regulators that promote EMT, apoptosis, resistance, angiogenesis, and tissue remodeling (37). Moreover, MMP9 has been associated with aggressive and metastatic breast cancer (38). In the invasive cancer tissue, we observed the activation of the Notch signaling pathway, a TNBC hallmark (39), through interaction with several ligands, for instance JAG2. Moreover, we identified the TNFSF10-TNFRSF10B interaction (Figure 4C) that commonly triggers apoptosis through caspases. The second apoptosis-triggering receptor TNFRSF10A was marginally expressed as well as TNFRSF10D, one of the two decoy receptors that modulate the apoptotic signal effectiveness. There are many resistance mechanisms downstream of the TNFRSF10A/B receptors to escape apoptosis (40), some of which were presumably active in the invasive cancer tissue.

**Figure 4.**
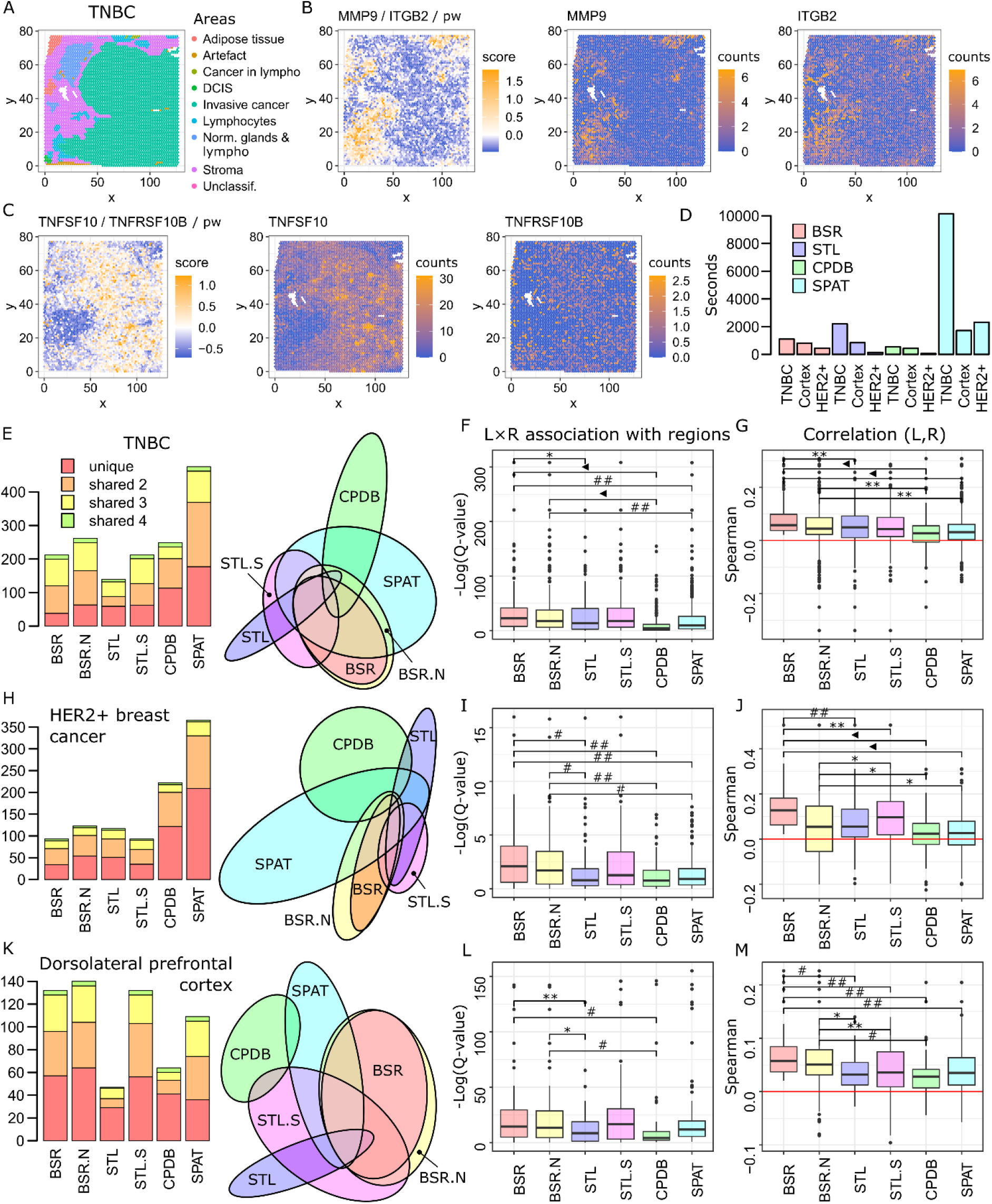
BulkSignalR for spatial transcriptomic data analysis. (**A**) Spatial organization of the TNBC tissue. (**B**) Example of a significant LRI associated with stroma. The left plot contains the gene signature scores of the (*L,R,pw*) triple where *pw* = interleukin 4 & 13 signaling (Reactome R-HAS-6785807). The middle and left plots show the ligand and receptor expression. (**C**) Example of LRI associated with cancer invasive tissue where *pw* = apoptosis (Reactome R-HSA-109581). (**D**) Computing times in seconds for the indicated tools using the indicated datasets. BSR=BulkSignalR, STL=stLearn, CPDB=CellPhoneDB, and SPAT=SpaTalk. (**E**) Number of significant LRIs in the TNBC dataset. BSR.N=BulkSignalR with negative L-R interactions (< −0.02), STL.S=stLearn second selection with a score threshold set to have the same number of LRIs as with BulkSignalR, STL=stLearn first selection. The color code indicates the number of tools that found one LRI (BSR and BSR.N as well as STL and STL.S count for one tool). The Venn diagram represents the overlap between tools and selections. (**F**) Statistical associations (Kruskal-Wallis followed by Benjamini-Hochberg multiple-hypothesis corrections) between tissue regions and concomitant expression of the ligand and the receptor captured by the product *L×R*. (**G**) Spearman correlation between ligand and receptor in the whole tissue. (**H**) Number of significant LRIs in the HER2^+^ breast cancer dataset. (**I,J**) Same as (F,G) for the HER2^+^ breast cancer dataset. (**K**) Number of significant LRIs in the dorsolateral prefrontal cortex dataset. (**L,M**) Same as (F,G) for the cortex dataset. *P < 0.05, **P < 0.01, #P < 1E-3, ##P < 1E-5, ◄P < 2.2E-16 (Wilcoxon 2-sided).

We obtained the plots in Figure 4A–C, the overview plot (Figure S17), statistical association with tissue regions (Figure S18), and a representation of LRI spatial pattern diversity (Figure S19) using BulkSignalR standard spatial functions.

### Comparison with existing spatial transcriptomic data analysis packages

We identified three recent or widely used tools that offer analysis at the multicellular resolution: stLearn (13), CellPhoneDB (10), and SpaTalk (14). We compared these tools with BulkSignalR using three human datasets: the previously used TNBC dataset (36), a HER2+ breast cancer dataset (41), and a dorsolateral prefrontal cortex dataset (42). To obtain comparable results, we employed LR*db* (our LRI database) with stLearn and SpaTalk. Despite our efforts, we could not replace the CellPhoneDB LRI database. Therefore, we used its native database that combines individual molecules and also complexes. When one interaction involved a complex (or two), we generated all possible pairwise LRIs. Then, we intersected this LRI list with LR*db* to be as close as possible to the other three tools. As for some LRIs, both molecules were given as ligands or receptors in LR*db*, we discarded these CellPhoneDB LRIs.

The next step was to apply comparable selection mechanisms. As not all tested tools implemented multiple hypothesis corrections, we used P-values for all tools including BulkSignalR. We imposed P values < 0.1% to remain close to the FDR < 1% we typically use with spatial data. stLearn required an adapted selection strategy. For each LRI, this tool combines CellPhoneDB and a local enrichment test at each spot *versus* its neighbors, and provides a P-value for each spot. We used a first selection mechanism that required at least 0.5% of the spots with P < 0.1%. This selection was rather sensitive to the number of imposed spots. Therefore, we used a second mechanism based on a score given by stLearn that is equal to the number of spots where a LRI was found with P < 5%. It was difficult to set a threshold for this score because multiple hypothesis correction would make most of these P-values non-significant. Hence, we simply took the same number of LRIs selected by BulkSignalR in decreasing order of this stLearn score.

CellPhoneDB and SpaTalk offer medium-resolution analysis, but they remain intrinsically optimized for single-cell or subcellular spatial resolution. Indeed, users must define a dominant cell type at each spot. In addition, CellPhoneDB requires the definition of the tissue regions (like in Figure 4A). They exploit this information to reconstitute cell type-specific transcriptomic profiles, restricted to each region for CellPhoneDB, or overall for SpaTalk. As SpaTalk reports LRIs by specifying the cell types, we selected all LRIs based on their P-values and eliminated redundant LRIs if they were significant in several cell type pairs. CellPhoneDB gives a P-value for each pair of cell types occurring in each region. We took the minimum P-value for each LRI and imposed a minimum P-value < 0.1%.

The TNBC, cortex, and HER2+ breast tumor datasets included 4,895, 3,636, and 306 spots, respectively. BulkSignalR, stLearn, and CellPhoneDB computing times scaled with the dataset size, while SpaTalk times were very long and difficult to explain (Figure 4D). CellPhoneDB was the fastest tool.

For the TNBC dataset, the four tools reported heterogeneous numbers of LRIs with limited overlap (Figure 4E). For CellPhoneDB and SpaTalk, we defined the dominant cell type at each spot according to the dataset authors who used single-cell data and a deconvolution software to determine them. For CellPhoneDB, we used the tumor regions defined by the authors and shown in Figure 4A. We obtained the largest number of LRIs with SpaTalk (twice as many as with the other tools), and the smallest number with stLearn (first selection). To relate the numbers of identified LRIs to their quality in the absence of a complete reference, we defined two objective quality criteria. First, we considered that the product of the ligand and the receptor, indicating co-presence at a spot, should display a statistical association with the tissue regions. Using this criterion, BulkSignalR performed significantly better than the other tools, with the exception of stLearn (second selection) (Figure 4F). This is remarkable because BulkSignalR does not exploit the knowledge of the tissue regions, unlike CellPhoneDB and SpaTalk. When we included negatively correlated ligands, BulkSignalR again performed better than CellPhoneDB and SpaTalk. CellPhoneDB was the worst. Second, we considered that the ligand and receptor abundances should be correlated in the whole sample. BulkSignalR output was significantly enriched in positive correlations compared with the other tools (Figure 4G). The other tools returned ~25% of LRIs with negative L-R correlations, which are more FP prone as reported above. Even when negative L-R correlations were allowed in BulkSignalR, the number of less reliable LRIs was significantly lower with BulkSignalR than with CellPhoneDB and SpaTalk.

Next, we compared the results obtained with the HER2+ breast tumor dataset. Tissue regions and dominant cell types were defined by the dataset authors. The number of significant LRIs identified by each tool was even more heterogeneous than for the TNBC dataset (Figure 4H). SpaTalk and CellPhoneDB found the largest number of interactions. Associations of the *L×R* product with tissue regions showed that BulkSignalR outperformed significantly SpaTalk and CellPhoneDB and also stLearn (first selection), but not stLearn (second selection). We obtained similar results when we allowed negative L-R correlations in BulkSignalR (Figure 4I). In terms of L-R correlations (Figure 4J), BulkSignalR outperformed all the other tools. CellPhoneDB and SpaTalk returned a very large number of negatively correlated LRIs, which might lead to substantial FP rates.

For the last comparison we used a dorsolateral prefrontal cortex dataset for which the authors defined regions, but no dominant cell types at each spot. As CellPhoneDB and SpaTalk compare remote spots anyway, we decided to apply them by using the regions as cell type definitions. stLearn (first selection) and CellPhoneDB returned the smallest lists of LRIs, while the other tools gave comparable numbers of LRIs (Figure 4K). Inter-tool heterogeneity remained substantial. Region association was better with BulkSignalR than CellPhoneDB and stLearn (first selection), but not compared with stLearn (second selection) and SpaTalk (Figure 4L). L-R correlations indicated that BulkSignalR performed better than the other tools.

### Application to colorectal cancer liver metastasis spatial data

To experimentally validate some LRIs identified by BulkSignalR, we generated a new ST dataset for four CRC-LM samples. Analysis of this dataset by BulkSignalR gave 173, 251, and 241 unique LRIs for the first three CRC-LM samples, respectively (FDR < 1%, L-R correlation > 0.02 in absolute value), and only 84 LRIs for the fourth sample (Table S1). As the obtained number of reads *per* spot was also smaller for the last sample (~30% less than the mean of the other three samples; data not shown), we excluded it. Application of clustering in Seurat combined with sample analysis by two pathologists allowed defining different regions in each sample (Figure 5A).

**Figure 5.**
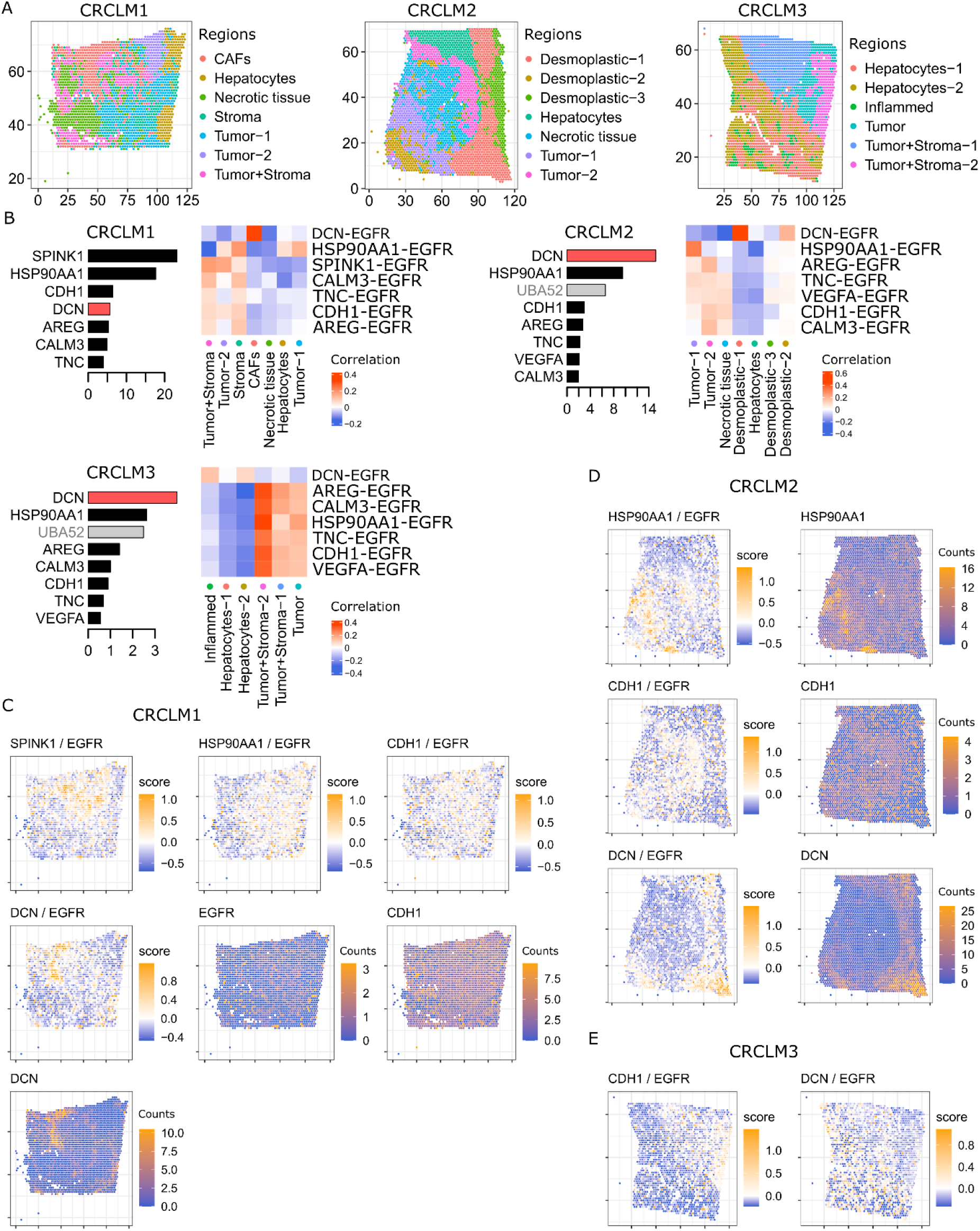
BulkSignalR analysis of a colorectal liver metastasis (CRC-LM) ST dataset. (**A**) Architecture of three CRC-LM samples. (**B**) Seven abundant EGFR ligands in the three CRC-LM. *DCN* is negatively correlated with EGFR. UBA52-EGFR was considered a dubious LRI and was ignored. (**C**) Spatial distribution of the four most abundant ligand-EGFR interactions (gene signature scores) in CRCLM1 and expression of *EGFR, CDH1*, and *DCN*. (**D**) Spatial distribution of the three most abundant ligand-EGFR interactions in CRCLM2. (**E**) Representative LRI spatial distributions in CRCLM3.

A complete analysis of the CRC-LM data with biological interpretation would obviously be out of the scope of this paper. We simply focused on LRIs with EGFR because its signaling is exacerbated in many tumors and EGFR is a clinical target in CRC-LM. EGFR has multiple ligands (21) and BulkSignalR identified 18, 19, and 15 of them in CRCLM1, CRCLM2, and CRCLM3, respectively. To assess their expression, we computed the 95^th^ percentile of their read counts in each sample. These revealed ligands that were strongly expressed in some area. To summarize, we represented the seven most abundant ligands in each CRC-LM sample in Figure 5B. We excluded the LRI UBA52-EGFR because ubiquitin A-52 residue ribosomal protein fusion product 1 (*UBA52*) does not seem to be secreted. The seven most expressed ligands displayed strong overlap in the three CRC-LM samples, despite the heterogeneous tumor architectures. One exception was serine protease inhibitor Kazal type 1 (*SPINK1*) that was the most expressed ligand in CRCLM1, but was not identified in CRCLM2/3. SPINK1 overexpression has been related to rare, fusion events in prostate cancer (43). It could be specific to CRCLM1 patient. Identification of the preferential expression of each ligand in each tumor region (see correlations in Figure 5B) showed that some ligands, *e.g*., decorin (*DCN*, and to a lower extent heat shock protein 90, *HSP90AA1)* were specific to some regions. DCN was always negatively correlated with EGFR, in line with its inhibitory action on EGFR signaling (44). DCN was also negatively correlated with cMET (*MET*) in all samples and the DCN-MET LRI has inhibitory effects (45). All other interactions involved positive L-R correlations.

Figure 5C shows the spatial configuration in CRCLM1 of the four most abundant ligand-EGFR interactions: SPINK1-, HAP90AA1-, E-cadherin (*CDH1*)-, and DCN-EGFR. The scores featured in the plots are the gene signature scores (Figure 1F) that combine ligand, receptor, and downstream pathway activity. We observed distinct areas of higher *versus* lower activity. We then plotted the expression of *EGFR, CDH1*, and *DCN* (Figure 5C). We observed that *EGFR* and CDH1 were almost ubiquitously expressed, whereas *DCN* expression was highest in cancer-associated fibroblast (CAF)-rich regions. This indicates that EGFR signaling could be potentially activated in each region of CRCLM1 through different ligands that are ubiquitously expressed or region-specific. The same analysis in CRCLM2 (Figure 5D) showed ligands with a specific localization (*DCN*), with intermediary localization (*HAP90AA1*), and with broad expression (*CDH1*). *EGFR* was almost ubiquitously expressed (data not shown). We obtained similar results for CRCLM3 (Figure 5E), although *DCN* expression was weaker and less localized due to the absence of desmoplastic reactions and CAF-rich regions in this sample.

We then experimentally validated the CDH1-EGFR and CDH1-cMET interactions in CRCLM1 and CRCLM2 to illustrate BulkSignalR capacity to identify interactions that are not necessarily sample region-specific. CDH1-EGFR and CDH1-cMET interactions have been described in many cell types; however, their relationship remains complex and context-dependent (46). The CDH1-MET interaction was not identified by BulkSignalR because it is not included in the LR*db* database. We assessed EGFR, cMET, CDH1 and phosphorylated ERK (pERK) abundance in the two CRC-LM samples by IF (Figures 6AB and S20AB). We used pERK as downstream reporter of EGFR and cMET signaling. In areas of pronounced CDH1 expression, EGFR membrane staining and pERK expression were less pronounced (Figures 6B and S20B). The pattern of cMET expression was similar to that of EGFR (stronger in CRCLM1 than CRCLM2). Reduction of EGFR intensity concomitantly with ERK phosphorylation was consistent with EGFR activation because upon phosphorylation, EGRF is rapidly internalized, thus decreasing its concentration at the membrane (47).

**Figure 6.**
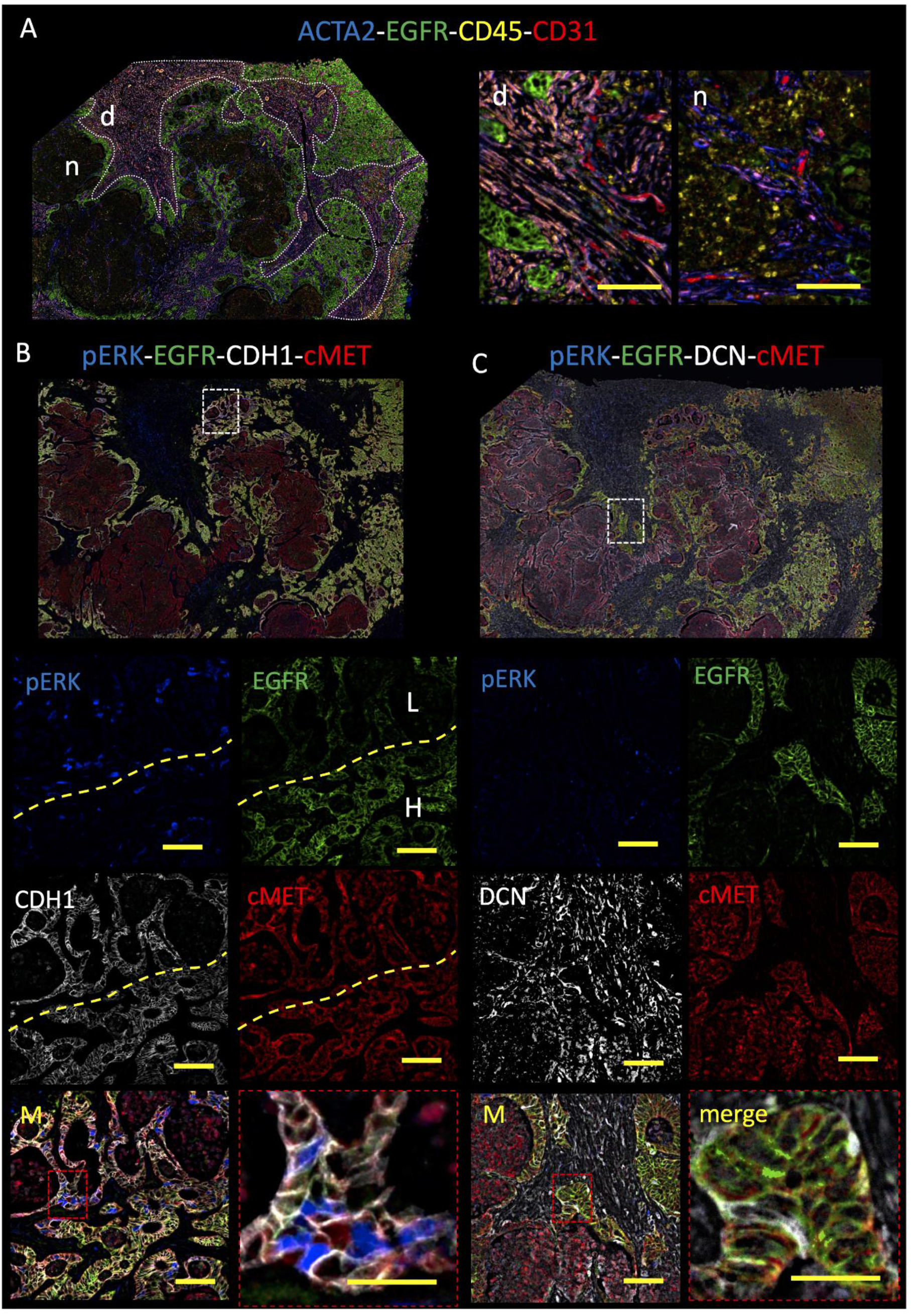
Validation by IF analysis of selected ligand-receptor interactions in CRCLM1. (**A**) (*left)* Structural overview of CRCLM1 with stromal areas (delineated in white) and EGFR^+^ cancer cells (green). ACTA2 is a CAF marker, CD45 is an immune cell marker, EGFR is cancer cell marker, and CD31 is an endothelial cell marker; d, desmoplastic reaction; n, necrosis. (*right)* Higher magnification view of desmoplastic and necrotic areas. (**B**) Analysis of CDH1-EGFR and CDH1-cMET interactions. Low-magnification view (100x) of quadruple staining (pERK, EGFR, CDH1, and cMET) in CRCLM1. The area of interest is highlighted and shown at higher magnification in the panels underneath. (**C**) Analysis of DCN-EGFR and DCN-cMET interactions. Same as in (B).

We then experimentally validated DCN-EGFR and DCN-MET interactions in CRCLM1/2 as examples of inhibitory LRIs. We consistently observed DCN expression in the desmoplastic area of the tumor, specifically in elongated stromal cells (Figures 6AC and S20AC). This was in agreement with previous reports that associated DCN expression with CAFs. We then investigated by IF the zones highlighted by the DCN-EGFR or DCN-MET LRI analysis. Activation of downstream signaling via pERK was clearly excluded in cancer cells, suggesting lower receptor tyrosine kinase activation in these cells. Overall, high stromal DCN expression was associated with lower receptor tyrosine kinase activation in both CRC-LMs.

## DISCUSSION

Invaluable transcriptomic and proteomic datasets are available and continue to be generated with a bulk methodology. Here, we showed that the R library BulkSignalR offers to researchers a solution to exploit these datasets to unravel cellular networks. BulkSignalR includes a rich set of functionalities, comparable to the best libraries for single-cell data analysis. The infrastructure that supports BulkSignalR computations to link LRIs and downstream pathways allows data analysis at both the pathway and individual LRI levels. There is an obvious parallel with enrichment analysis of gene sets *versus* the analysis of individual differentially expressed genes. This infrastructure also allows network visualization for relating LRIs to target genes. Although bulk data do not directly convey information about the specific transcriptomes of individual cell populations, we propose a simple machine learning model that can infer what populations are likely to participate in each LRI.

By comparing the LRIs inferred by BulkSignalR from several bulk datasets and the corresponding single-cell datasets, we found that the single-cell data analysis typically identified 2-3 times more LRIs. We also found that LRIs identified in bulk data only tended to involve low-abundance ligands and receptors, which is in line with the general higher sensitivity of bulk technologies. Therefore, single-cell analysis is the approach of choice to map cellular networks when few representative samples are sufficient. In the other cases, bulk approaches can be used for cellular network inference, but with reduced details. The two formats, bulk and single-cell, could be eventually combined as well as data analysis methods in large studies.

ST is rapidly developing and a frequent setting consists in working at multicellular resolution. Specifically, a tissue section is probed at multiple locations (on a grid), and each spot has a size that results in the concomitant analysis of more than one cell. For instance, in the very popular Visium^™^ system, a spot contains between 3 and 30 cells, in function of the tissue and cell types. Therefore, the transcriptome data acquired at each spot are bulk by nature and BulkSignalR is suitable for analyzing such ST data. By reviewing the literature, we discovered that most existing ST software tools have been developed for single-cell or subcellular resolution data. Some nevertheless claim to be compatible with Visium-type data, for instance CellPhoneDB (10), SpaTalk (14), and stLearn (13). In stLearn, an initial CellPhoneDB application is combined with a local posterior analysis to compare a spot with its neighbors. Conversely, CellPhoneDB alone and SpaTalk consider LRIs between spots that may be far from each other. Although some LRIs have a longer range, many occur at a short range or only at cell contacts. We think that this renders the output of these tools difficult to interpret. Nevertheless, we compared CellPhoneDB, SpaTalk, and stLearn with BulkSignalR using three ST datasets. We found that BulkSignalR inferred an average number of LRIs who were significantly more reliable according to two neutral quality criteria. The number of LRIs found by stLearn, CellPhoneDB and SpaTalk varied substantially, and SpaTalk computing times were very long (up to hours on four processors). We also observed limited overlap between the four tools for each dataset.

Using a new Visium^™^ dataset that included four CRC-LM samples, we used two of these samples to experimentally validate selected inferences obtained with BulkSignalR. We confirmed the CDH1-EGFR, CDH1-cMET, DCN-EGFR, and DCN-cMET interactions in both samples by IF. The activation of receptors, and especially receptor tyrosine kinases, such as EGFR, is complex and involves multiple agonist and antagonist ligands, as illustrated by our analysis that unraveled several, spatially-dependent ligands besides CDH1 and *DCN*. The complete analysis of the net results of all these interactions goes beyond the scope of this article, but the presented validations showed the potential of our ST data analysis pipeline.

The analysis of spatial data did not require any modification of BulkSignalR, only the adaptation of few parameters. However, we added dedicated graphical functions. The BulkSignalR R library was designed to be easy to use by scientists with basic knowledge of R. Few functions are sufficient to prepare a dataset, infer the LRIs, and generate informative plots.

## AVAILABILITY

BulkSignalR is available with documentation from https://github.com/jcolinge/BulkSignalR.

## ACCESSION NUMBERS

The CRC-LM spatial transcriptomics data will be available from GEO upon publication. The other data were made public by their authors.

## SUPPLEMENTARY DATA

Supplementary Data (Tables S1 and S2, Supplementary Information) are available at NAR online.

## ACKNOWLEDGEMENT

The authors acknowledge the experimental help of Mr. Cyril Degletagne, Cancer Genomics Platform (Centre Léon Bérard, Cancer Research Center of Lyon, France) for NGS library preparation and sequencing. The authors are thankful to Dr. Virginie Georget (Montpellier Imaging Platform, Biocampus, Montpellier) for the help with IF data acquisition.

