## Supplementary Information for "Inferring ligand-receptor cellular networks from bulk and spatial transcriptomic datasets with BulkSignalR"

### Supplementary Figures

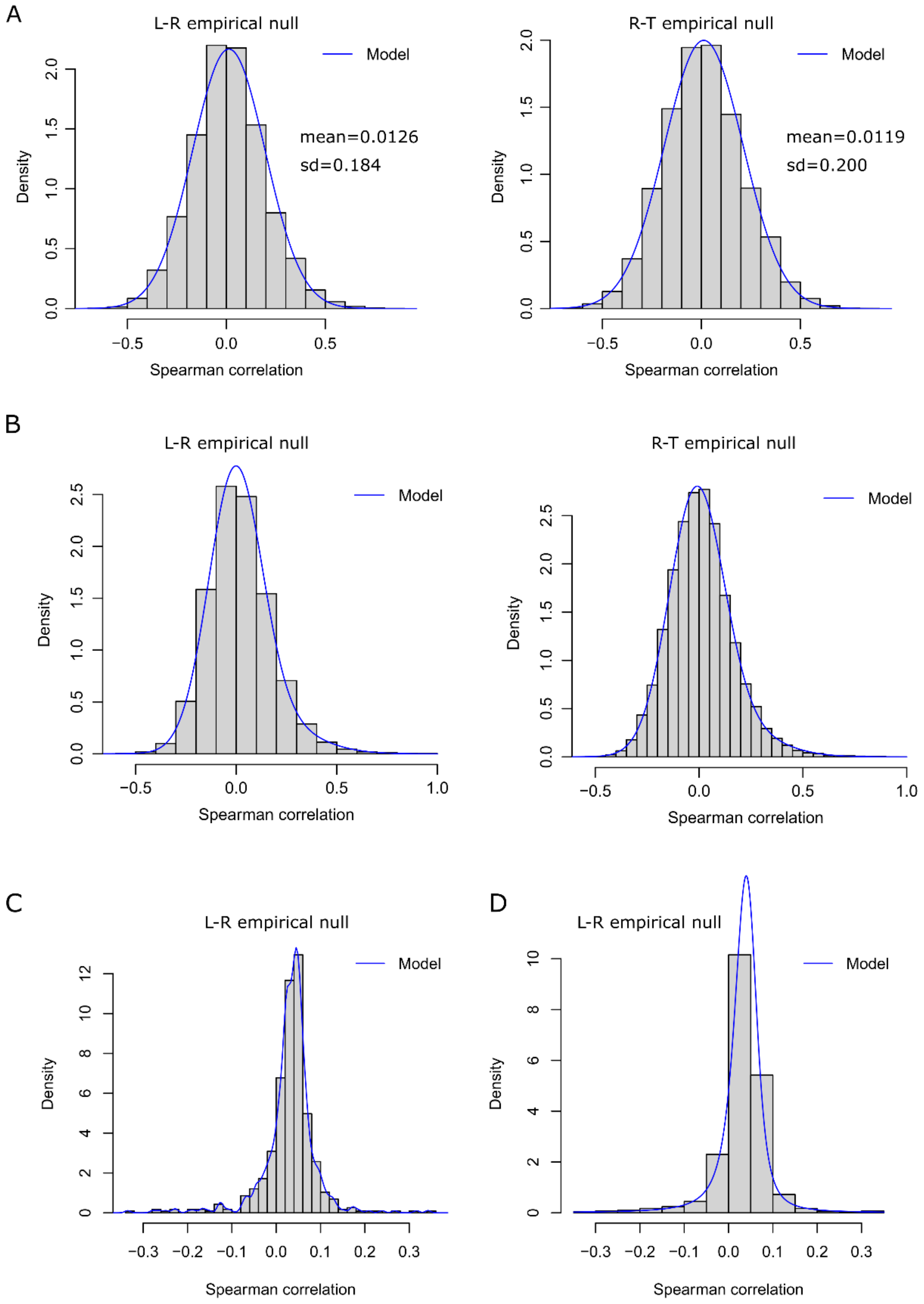


**Figure S1.** Representative examples of null distributions for RNA-seq data. (**A**) The SDC (1, 2) dataset fit with a censored normal distribution. We note the close parameters for the L-R and R-T null distributions. (**B**) The sarcoma Affymetrix microarray dataset (3) fit by a censored mixture of two normal distributions. (**C**) Gaussian kernel-based empirical model on the human dorsolateral prefrontal cortex dataset. (**D**) Censored stable model on the same dataset.

| L | R | pw.id | pw.name | rank | len | qval |
|---|---|---|---|---|---|---|
| BTLA | CD247 | GO:0050852 | T cell receptor signaling pathway | 36 | 56 | 1.3E-53 |
| B2M | CD247 | GO:0050852 | T cell receptor signaling pathway | 36 | 56 | 5.5E-51 |
| BTLA | CD247 | GO:0002250 | adaptive immune response | 26 | 40 | 5.1E-49 |
| B2M | CD247 | GO:0002250 | adaptive immune response | 26 | 40 | 3.2E-46 |
| IL16 | CD4 | R-HSA-202403 | TCR signaling | 32 | 50 | 4.0E-46 |
| IL16 | CD4 | R-HSA-388841 | Costimulation by the CD28 family | 32 | 50 | 1.7E-45 |
| B2M | CD3D | R-HSA-202403 | TCR signaling | 32 | 50 | 2.1E-43 |
| CXCL12 | CD4 | R-HSA-202403 | TCR signaling | 32 | 50 | 2.1E-43 |
| HLA-B | CD3D | R-HSA-202403 | TCR signaling | 32 | 50 | 2.1E-43 |
| HLA-G | CD4 | R-HSA-202403 | TCR signaling | 32 | 50 | 2.1E-43 |
| HLA-A | CD3D | R-HSA-202403 | TCR signaling | 32 | 50 | 3.1E-43 |
| CXCL12 | CD4 | R-HSA-388841 | Costimulation by the CD28 family | 32 | 50 | 8.1E-43 |
| HLA-G | CD4 | R-HSA-388841 | Costimulation by the CD28 family | 32 | 50 | 8.1E-43 |
| LILRB4 | LAIR1 | GO:0002250 | adaptive immune response | 24 | 38 | 2.1E-37 |
| CXCL12 | CD4 | R-HSA-202424 | Downstream TCR signaling | 25 | 39 | 2.5E-33 |
| NPPA | NPR1 | R-HSA-397014 | Muscle contraction | 103 | 159 | 8.1E-33 |
| LTA | TNFRSF1B | R-HSA-6783783 | Interleukin-10 signaling | 26 | 40 | 5.1E-32 |
| LTA | TNFRSF1B | R-HSA-6785807 | Interleukin-4 and Interleukin-13 signaling | 54 | 84 | 1.6E-31 |
| OMG | TNFRSF1B | R-HSA-6783783 | Interleukin-10 signaling | 26 | 40 | 3.7E-31 |
| IL10 | IL10RA | R-HSA-6783783 | Interleukin-10 signaling | 26 | 41 | 7.8E-31 |
| OMG | TNFRSF1B | R-HSA-6785807 | Interleukin-4 and Interleukin-13 signaling | 54 | 84 | 1.1E-30 |
| TNF | TNFRSF1B | R-HSA-6783783 | Interleukin-10 signaling | 26 | 40 | 2.6E-30 |
| TNF | TNFRSF1B | R-HSA-6785807 | Interleukin-4 and Interleukin-13 signaling | 54 | 84 | 8.1E-30 |
| … | … | … | … | … | … | … |

**Figure S2.** Typical contents of the tabular representation of (ligand, receptor, pathway) triples with statistical significance. We note redundancy in the table due to inherent redundant Reactome pathways and GO biological processes. Reduction operators enable alleviating this phenomenon. The full table was 4,912 rows long. Len stands for the number of target genes in the pathway and rank was the position of the percentile used for order statistics, *e.g.*, for the first example, rank 36 is 65$^{th}$ percentile (from 56 sorted target gene correlations).

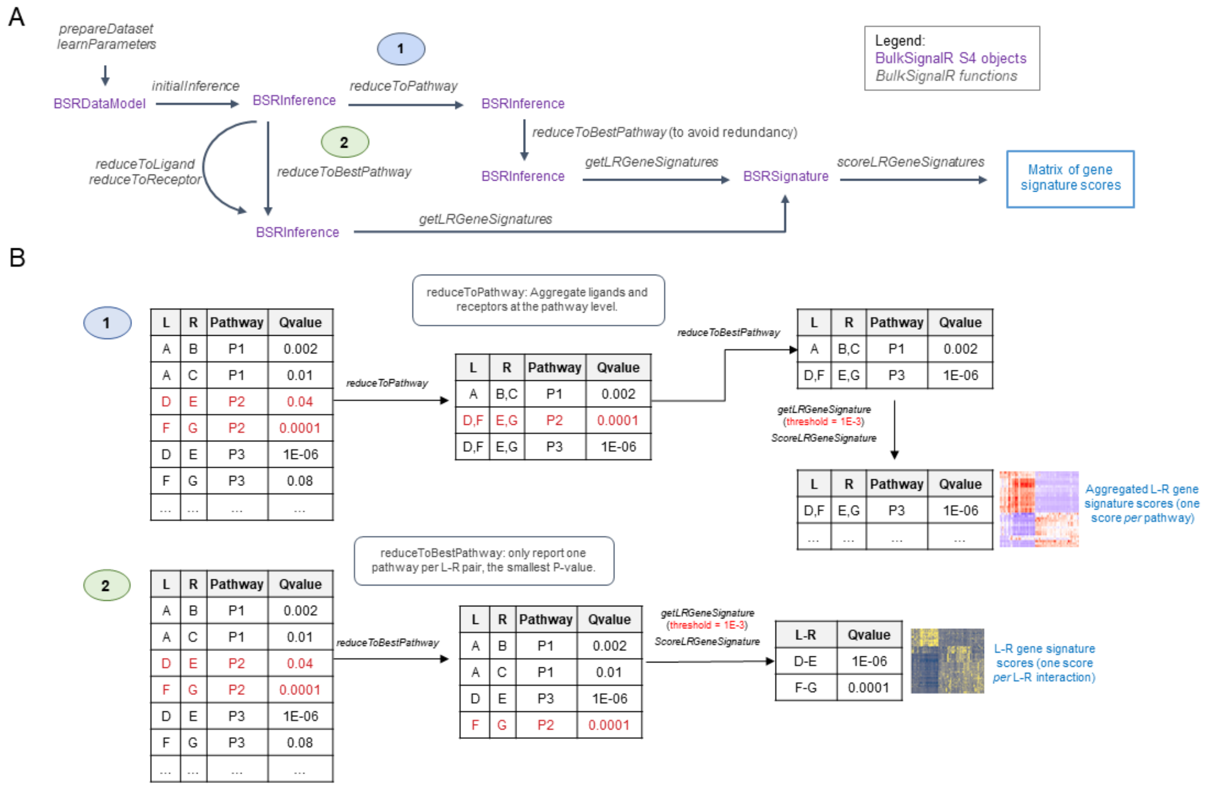


**Figure S3.** The different reduction operations available in BulkSignalR. (**A**) Typical alternative combinations of reducing complexity, computing gene signatures, and scoring the latter. (**B**) Differences between reducing to pathways (1, top) and reducing to the best pathway (2, bottom). In every case, gene signatures are comprised of the ligand(s), receptor(s), and downstream pathway target genes. The scores include the contributions (z-scores) of each of these molecules.

A

**Pancreas**

| BulkSignalR | | Smartseq coverage ( M = 1,119 ) | |
|---|---|---|---|
| FDR | N | % | log10(P) |
| 1E-03 | 400 | 57.0% | -22 |
| 1E-04 | 357 | 59.1% | -23 |
| 1E-05 | 322 | 59.9% | -22 |
| 1E-06 | 263 | 61.6% | -20 |

B

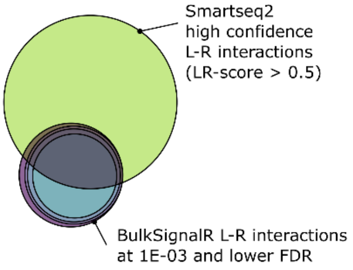


C

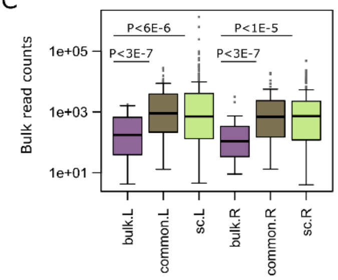


D

**Liver**

| BulkSignalR | | Chromium coverage (M = 1,860 ) | | mCEL-Seq2 coverage ( M = 2,368 ) | |
|---|---|---|---|---|---|
| FDR | N | % | log10(P) | % | log10(P) |
| 1E-03 | 297 | 85.9% | -28 | 95.3% | -25 |
| 1E-04 | 192 | 92.2% | -28 | 95.8% | -17 |
| 1E-05 | 132 | 93.2% | -20 | 97.0% | -13 |
| 1E-06 | 66 | 92.4% | -10 | 93.9% | -5 |

E

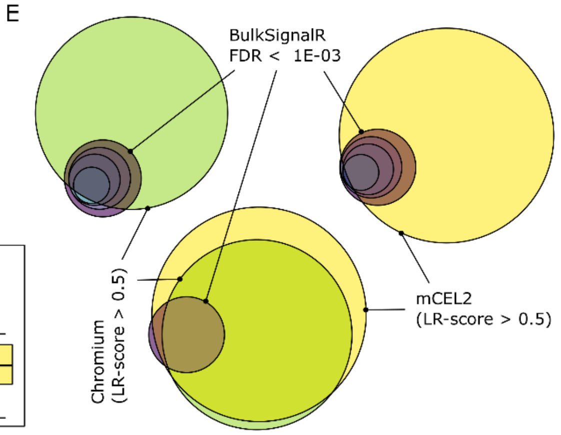


F

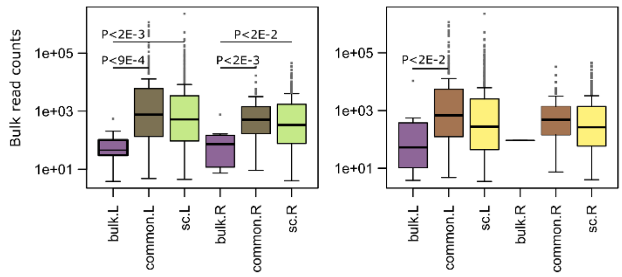


G

**Brain cortex**

| BulkSignalR | | Chromium coverage ( M = 2,344 ) | | Smartseq coverage ( M = 2,286 ) | |
|---|---|---|---|---|---|
| FDR | N | % | log10(P) | % | log10(P) |
| 1E-03 | 522 | 89.3% | -24 | 93.1% | -43 |
| 1E-04 | 477 | 89.3% | -22 | 93.5% | -40 |
| 1E-05 | 436 | 89.7% | -21 | 93.6% | -37 |
| 1E-06 | 355 | 92.1% | -22 | 92.7% | -27 |

H

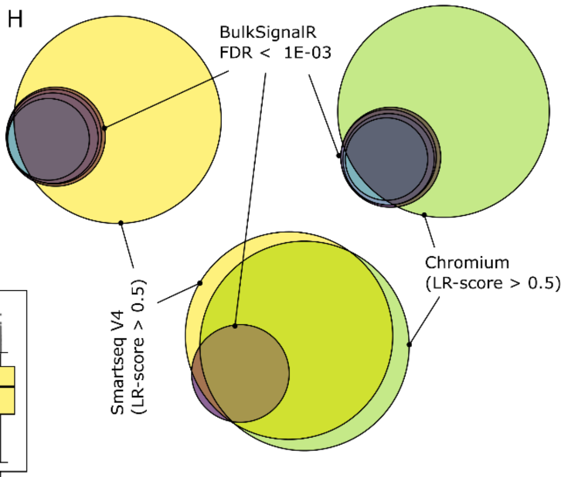


I

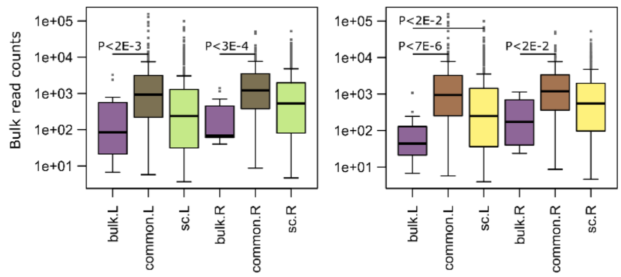


**Figure S4.** Comparison with matching single-cell datasets. (**A**) Table with the number of unique L-R interactions below a certain false discovery rate (FDR) or Q-value found in GTEx pancreas bulk transcriptomes by BulkSignalR. These numbers were compared with the number (M = 1,119) of high-confidence L-R interactions found in a Smartseq pancreas dataset (4) with SingleCellSignalR (see Supplementary Methods). The right-hand part of the table shows how much of the bulk-found interactions were covered by the single-cell-found interactions, and the P-value associated with such a coverage (hypergeometric test). (**B**) Venn diagram representing the overlap for the various FDR thresholds. (**C**) Comparison of the read counts in the bulk GTEx dataset of the ligands and receptors found in the L-R interactions unique to the bulk, common, and unique to single-cell (BulkSignalR FDR $< 10^{-3}$). We note that ligands and receptors involved in bulk-specific L-R interactions are less expressed. (**D**) Same as (A) with GTEx liver bulk transcriptomes, and a Chromium (5) and a mCEL-Seq2 (6) single-cell datasets. (**E**) Same as (B) but with two single-cell datasets. (**F**) Same as (C). (**G**) Same as (A) with GTEx brain frontal cortex transcriptomes, and a Chromium (7) and a Smartseq (8) single-cell datasets. (**H**) Same as (E). (**I**) Same as (F).

```r
library(BulkSignalR)

# read expression data
counts <- read.csv(...)

# obtain a BulkSignaR Data Model object & learn statistical parameters
bsrdm <- prepareDataset(counts)
bsrdm <- learnParameters(bsrdm)

# infer the L-R interactions as a BulkSignalR Inference object
bsrinf <- initialInference(bsrdm)

# interactions can be obtained as a data.frame with an accessor
inter <- LRinter(bsrinf)  # see Fig. S4 for the content of this data.frame

# a list of vectors containing the target genes of each pathway
# can be obtained via another accessor, same for corresp. correlations
tgenes <- tGenes(bsrinf)
tgcorr <- tgCorr(bsrinf)

# computation of gene signatures as well as their scoring
bsrsig <- getLRGeneSignatures(bsrinf, qval.thres=1e-3)
scores <- scoreLRGeneSignatures(bsrdm, bsrsig)
simpleHeatmap(scores)

# reductions return new BulSignalR Inference objects
bsrinf.red <- reduceToBestPathway(bsrinf)
head(LRinter(bsrinf.red))
bsrsig.red <- getLRGeneSignatures(bsrinf.red, qval.thres=1e-3)
scores.red <- scoreLRGeneSignatures(bsrdm, bsrsig.red)
simpleHeatmap(scores.red)

# reduction to the ligand
bsrinf.redL <- reduceToLigand(bsrinf)

# reduction to the receptor
bsrinf.redR <- reduceToReceptor(bsrinf)

# reduction to pathway
bsrinf.redP <- reduceToBestPathway(reduceToPathway(bsrinf))
```

**Figure S5.** Typical sequence of R commands using the BulkSignalR library.

**Figure S6 (next page).** Illustration of BulkSignalR network representation capabilities. (**A**) L-R interactions can be represented (igraph R package) and plotted as graphs. Here we show L-R interactions involved in the six brain pathways that were highlighted in Figure 2B. (**B**) BulkSignalR tabular output for the six chosen pathways. (**C**) Chord diagram representation. (**D**) Bubble plot representation. (**E**) L-R interactions extended down to the six chosen pathway target genes through shortest paths. (**F**) While BulkSignalR network functions can return igraph objects that can be plotted in R directly as in panels A and E, igraph also offers functions to export graphs in standard file formats such as graphML. This enables importing networks into other software packages to plot and analyze interactions such as yEd (www.yworks.com) or Cytoscape (9).

A

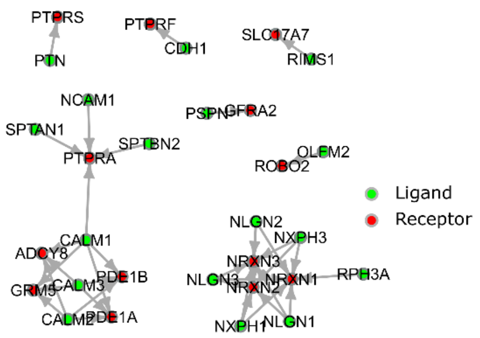


B

| L | R | pw.id | pw.name | qval |
|---|---|---|---|---|
| CALM1 | PTPRA | R-HSA-375165 | NCAM signaling for neurite out-growth | 5.69E-05 |
| NCAM1 | PTPRA | R-HSA-375165 | NCAM signaling for neurite out-growth | 2.57E-04 |
| PSPN | GFRA2 | R-HSA-375165 | NCAM signaling for neurite out-growth | 4.31E-05 |
| SPTAN1 | PTPRA | R-HSA-375165 | NCAM signaling for neurite out-growth | 1.04E-04 |
| SPTBN2 | PTPRA | R-HSA-375165 | NCAM signaling for neurite out-growth | 3.85E-05 |
| CALM1 | GRM5 | R-HSA-6794361 | Neurexins and neuroligins | 1.34E-19 |
| CALM2 | GRM5 | R-HSA-6794361 | Neurexins and neuroligins | 1.19E-19 |
| CALM3 | GRM5 | R-HSA-6794361 | Neurexins and neuroligins | 1.53E-19 |
| NLGN1 | NRXN1 | R-HSA-6794361 | Neurexins and neuroligins | 3.81E-22 |
| NLGN1 | NRXN2 | R-HSA-6794361 | Neurexins and neuroligins | 1.94E-21 |
| NLGN1 | NRXN3 | R-HSA-6794361 | Neurexins and neuroligins | 5.78E-22 |
| NLGN2 | NRXN1 | R-HSA-6794361 | Neurexins and neuroligins | 7.72E-23 |
| NLGN2 | NRXN2 | R-HSA-6794361 | Neurexins and neuroligins | 2.65E-22 |
| NLGN2 | NRXN3 | R-HSA-6794361 | Neurexins and neuroligins | 8.08E-23 |
| NLGN3 | NRXN1 | R-HSA-6794361 | Neurexins and neuroligins | 4.14E-23 |
| NLGN3 | NRXN2 | R-HSA-6794361 | Neurexins and neuroligins | 2.16E-22 |
| NLGN3 | NRXN3 | R-HSA-6794361 | Neurexins and neuroligins | 5.71E-23 |
| NXPH1 | NRXN1 | R-HSA-6794361 | Neurexins and neuroligins | 3.05E-23 |
| NXPH1 | NRXN2 | R-HSA-6794361 | Neurexins and neuroligins | 2.64E-22 |
| NXPH1 | NRXN3 | R-HSA-6794361 | Neurexins and neuroligins | 2.91E-23 |
| NXPH3 | NRXN1 | R-HSA-6794361 | Neurexins and neuroligins | 1.77E-22 |
| NXPH3 | NRXN2 | R-HSA-6794361 | Neurexins and neuroligins | 1.21E-21 |
| NXPH3 | NRXN3 | R-HSA-6794361 | Neurexins and neuroligins | 4.24E-22 |
| RPH3A | NRXN1 | R-HSA-6794361 | Neurexins and neuroligins | 4.01E-23 |
| RIMS1 | SLC17A7 | R-HSA-112310 | Neurotransmitter release cycle | 6.11E-16 |
| CALM1 | ADCY8 | R-HSA-111885 | Opioid Signalling | 4.13E-10 |
| CALM1 | PDE1A | R-HSA-111885 | Opioid Signalling | 1.62E-10 |
| CALM1 | PDE1B | R-HSA-111885 | Opioid Signalling | 2.28E-08 |
| CALM2 | ADCY8 | R-HSA-111885 | Opioid Signalling | 3.39E-10 |
| CALM2 | PDE1A | R-HSA-111885 | Opioid Signalling | 3.55E-10 |
| CALM2 | PDE1B | R-HSA-111885 | Opioid Signalling | 3.07E-08 |
| CALM3 | ADCY8 | R-HSA-111885 | Opioid Signalling | 3.62E-10 |
| CALM3 | PDE1A | R-HSA-111885 | Opioid Signalling | 4.26E-10 |
| CALM3 | PDE1B | R-HSA-111885 | Opioid Signalling | 2.37E-08 |
| OLFM2 | ROBO2 | R-HSA-9010642 | ROBO receptors bind AKAP5 | 1.59E-04 |
| CDH1 | PTPRF | R-HSA-8849932 | Synaptic adhesion-like molecules | 9.09E-04 |
| PTN | PTPRS | R-HSA-8849932 | Synaptic adhesion-like molecules | 2.88E-04 |

C

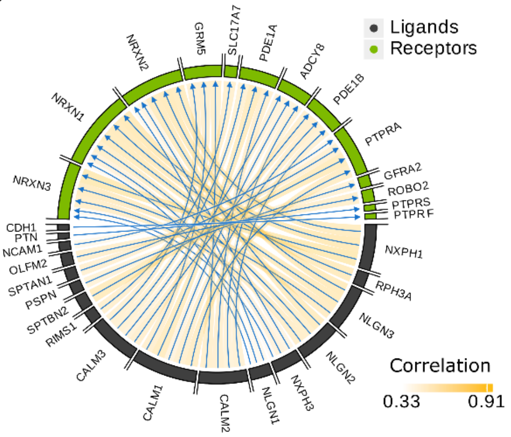


D

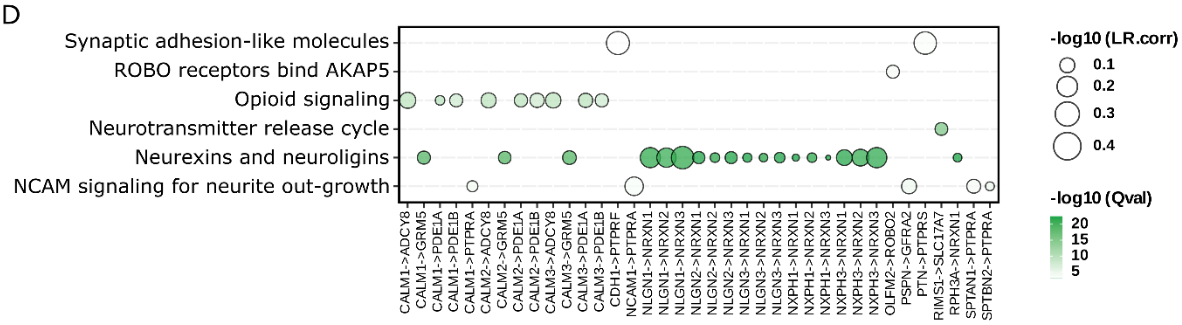


E

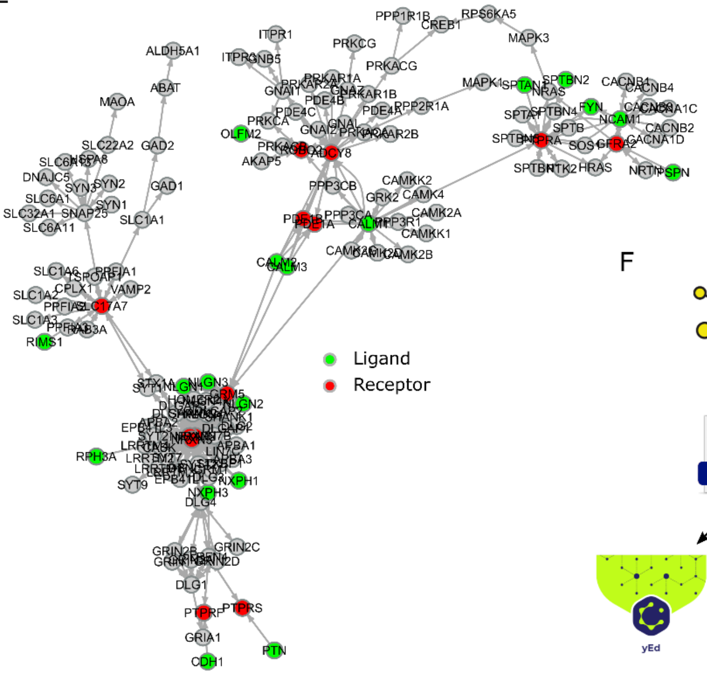


F

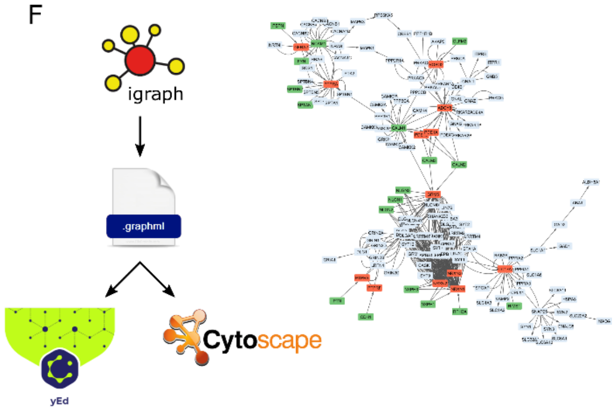

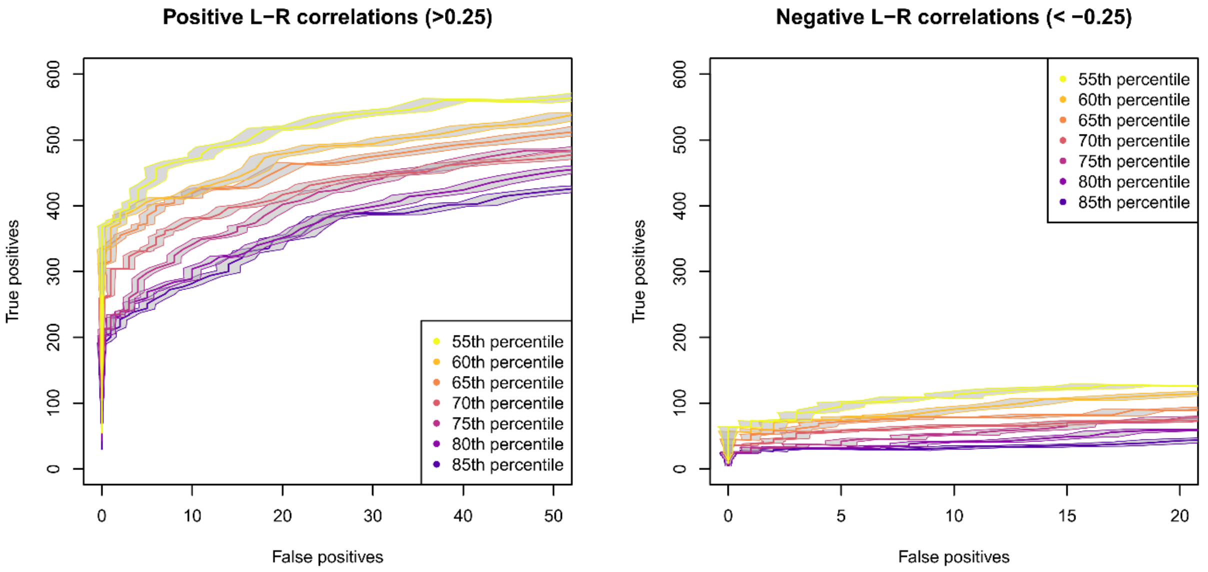


**Figure S7.** Pseudo-ROC curves for positive and negative L-R correlations in the SDC dataset. The curves for positive (> 0.25) L-R correlations (left) reproduce what is observed when only the positive correlations are allowed (Figure 1D). The curves for negative (< -0.25) L-R correlations are much worse (right) with roughly five times less TPs for a given number of FPs.

### Relating a set of genes to ligands

A single-cell tool, NicheNet (10), proposed to exploit an integrated molecular interaction network, including LRIs and a notion of target genes, to relate user-chosen sets of genes to ligands that should drive their expression. They demonstrated this functionality using the 100-gene signature proposed by Puram (11) for a partial EMT (p-EMT) cancer cell transcriptional program taking place at the invasive front of some head and neck squamous cell carcinoma (HNSC). Working at single-cell resolution, Puram *et al.* suggested that ligands produced by cancer-associated fibroblasts (CAFs) proximal to the invasive front might interact with cancer cell receptors to activate p-EMT. Detailed validation work was conducted on the influence of TGF-β3.

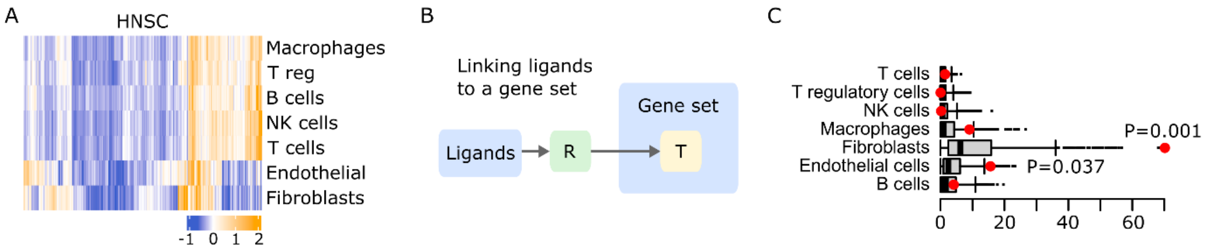


**Figure S8.** (**A**) Scoring of common cell type signatures across 500 TCGA HNSC transcriptomes. (**B**) Principle of relating a ligand to genes from a given gene set through BulkSignalR L-R-pathway triples. (**C**) Empirical significance analysis of the associations between the LRIs involving the retrieved ligands for p-EMT and cell types in (E).

Obviously, there is conceptual proximity between NicheNet integrated reference network and how we link ligands down to receptor-pathway target genes. We thus decided to use BulkSignalR inferences to provide a functionality close to NicheNet, but for bulk data. We processed TCGA primary HNSC data that were comprised of 500 transcriptomes with BulkSignalR default parameters. Common cell type signatures were again employed to score their abundance over the whole cohort (Figure S8A). Individual LRI inferences, *i.e.*, $(L, R, pw)$ triples, were imposed FDR < 0.1% and at least 2 p-EMT genes as targets of $pw$ (Figure S8B). This selected 80 ligands that were not already in p-EMT, including *TGFB1* and *TGFB3*. We then computed associations with cell types as explained in the main text, which gave a total association score for each cell type by summing the weights in each linear model (red dots in Figure S8C). We repeated the overall procedure 1,000 times for randomly chosen gene sets with similar intersection size with BulkSignalR pathway targets (45 out of 100 p-EMT genes were in our targets). Accordingly, we derived empirical estimates of statistical significance and found CAFs to be highly significant and endothelial cells marginally (Figure S8C). This specific example shows the potential to extract information relating gene regulation and signal-emitting cell populations from bulk data. The overall procedure is implemented in BulkSignalR functions for users to apply it to datasets and gene sets of their choice.

### Autocrine interactions of breast cancer cell lines in culture conditions

Transcriptomes of 90 breast cancer cell lines from DepMap project (12) were analyzed at the pathway level (FDR < 0.1%), full tabular output in Table S2. The resulting signature scores separated the major basal-like and luminal molecular subtypes (Figure S9), which arise from different cells of origin and show distinct biological characteristics and invasive potential (13, 14). HER2+ tumor cell lines appeared mixed within the luminal class, which is consistent with later integrative, molecular-based stratifications that classified HER2+ and some Luminal B tumors as a specific subgroup (15, 16).

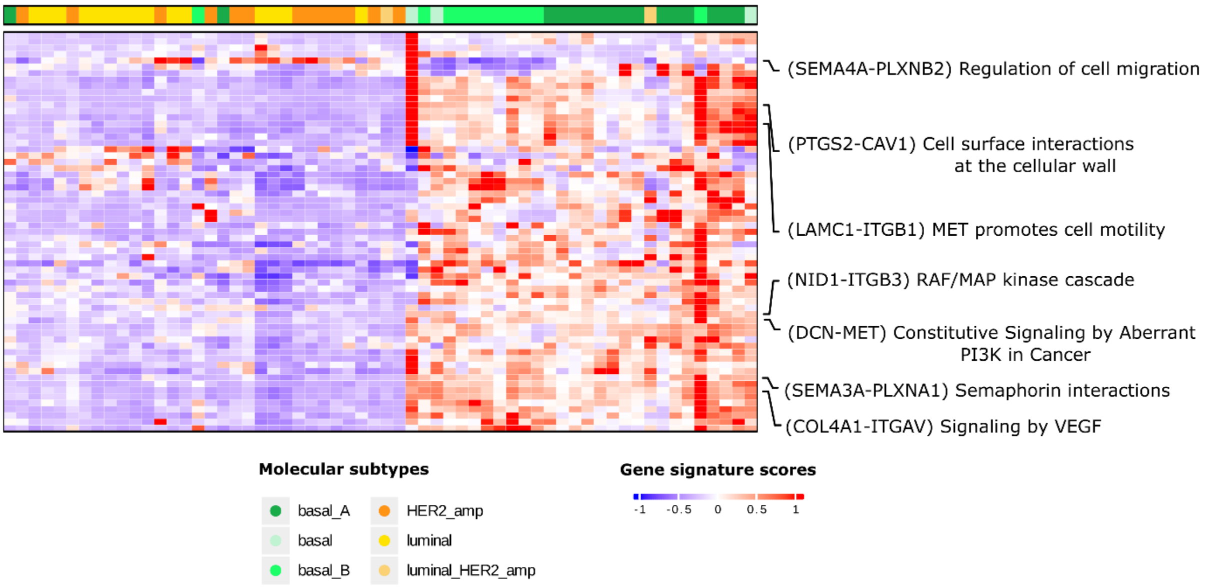


**Figure S9.** Pathway activity across 90 breast invasive carcinoma cell lines. Some illustrative pathways along with one representative L-R interaction involved are highlighted. Molecular subtypes were obtained from DepMap.22Q2 portal.

As expected, basal cell lines featured higher gene signature scores for pathways related to EMT and augmented cell motility (see also Figure S10A). Logically, regulation of cell migration was essentially the only downregulated pathway in basal cells (Figure S10B).

The PTGS2-CAV1 L-R interaction relates to Cell surface interactions at vascular wall pathway, which displayed stronger activity in basal cells (Figure S10C) and allows white cells to leave circulation and enter the tumor by extravasation. Caveolin-1 (gene *CAV1)* is a multifunctional scaffolding protein with multiple binding partners that associates with cell surface caveolae, and is considered an important player in the progression of breast carcinoma. It modulates several biological functions such as cell proliferation, angiogenesis, migration and therapeutic resistance (17). *CAV1* expression has been found increased in breast tumors with a basal-like phenotype (18). Furthermore, PI3K and MAP kinase signaling, which play important roles in the growth and metastatic potential of breast cancer (19–21), appeared through the DCN-MET and NID1-ITGB3 interactions, respectively. Their higher activity in basal cancer cells (Figure S10DE) is consistent with previous findings (22, 23).

Interestingly, we also noted a difference between breast cancer subtypes for semaphorin interactions (Figure S10F). Semaphorins are starting to be considered as potential clinical biomarkers and therapeutic targets in breast cancer (24).

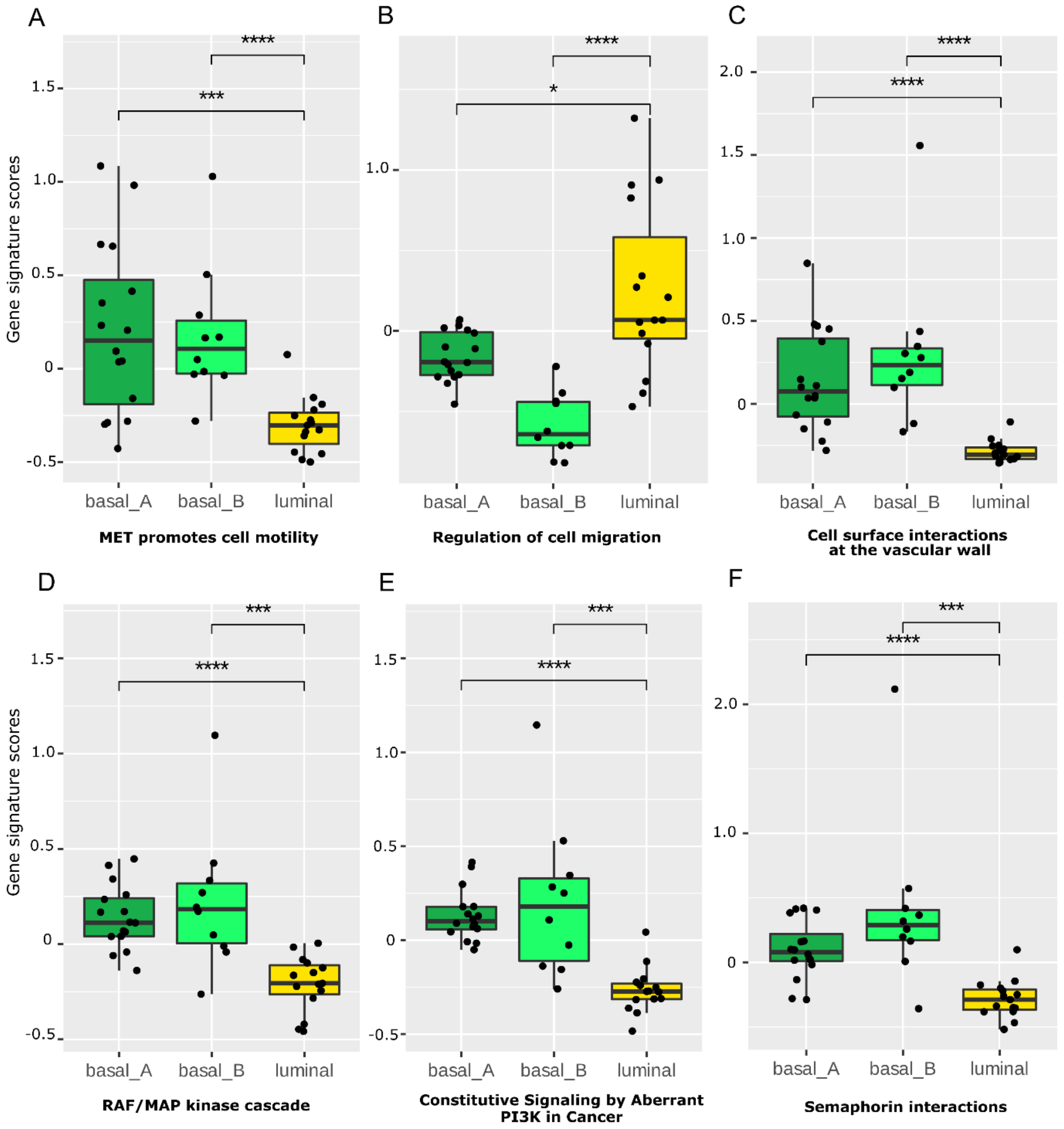


**Figure S10.** Differential activity of chosen pathways. (Wilcoxon test, *** P < 0.001, **** P < 0.0001).

In breast cancer, extracellular matrix (ECM) components such as collagens, *i.e.*, COL4A1 here, are significantly deregulated and are known to influence cell growth, migration and angiogenesis by interacting the cell surfaces through multiple integrin receptors, forming a highly dynamic molecular network (25). In Figure S11, we highlight the intricacy of COL4A1 interactions linking this protein to several receptors and downstream pathways thus showing BulkSignalR potential to infer such complex, but highly relevant networks.

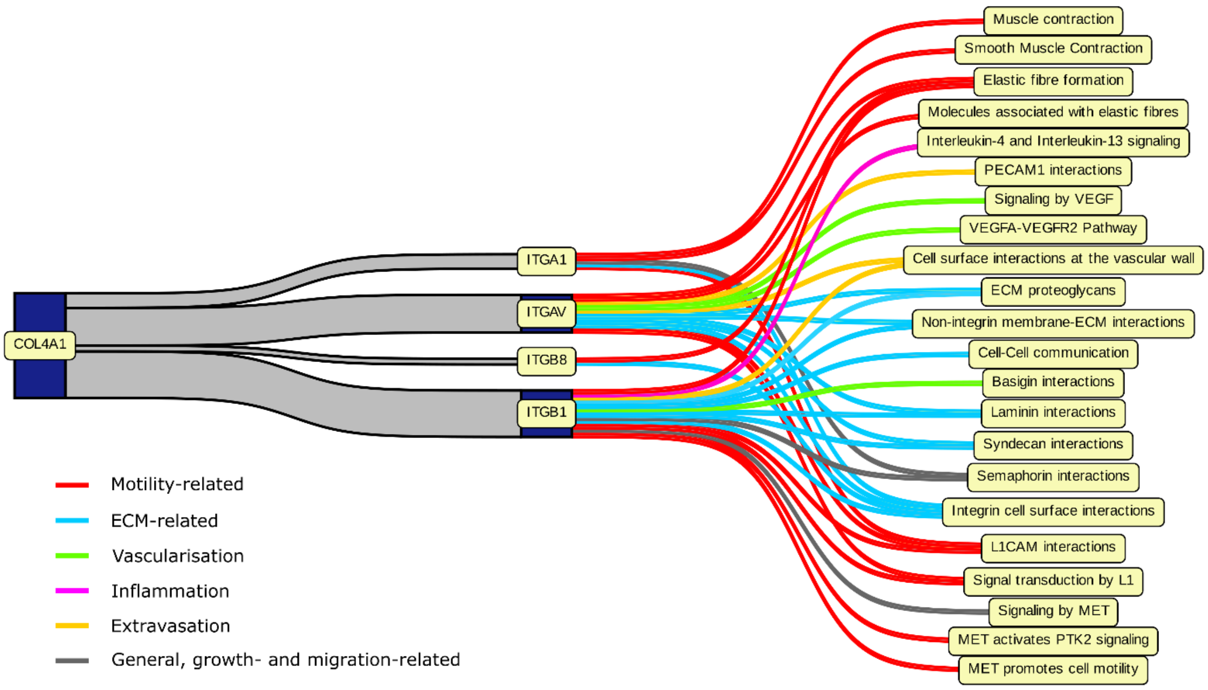


**Figure S11.** Sankey (or lava) plot showing how COLA41 relates to several pathways through different receptors.

To corroborate our results with an independent dataset as well as to showcase the application of BulkSignalR to a different data type, we downloaded expression proteomics data of 30 breast cancer cell lines from DepMap 22Q2 Portal (Supplementary Methods). In agreement with transcriptomics results, a clear distinction between the luminal and basal phenotypes was obtained (Fig. S12A, Table S2). Pathways related to the ECM and cell-cell/cell-matrix interactions dominated and were increased in the basal subtype. This highlighted again the critical importance of ECM remodeling in breast cancer pathogenesis (26). The protein L-R interactions matched the gene L-R interactions significantly (Fig. S12B). Although involving different ligand and receptor molecules, some of these pathways showed a clear concordance with pathways found in the transcriptomics dataset such as Cell surface interactions at the vascular wall (Fig. S12CD).

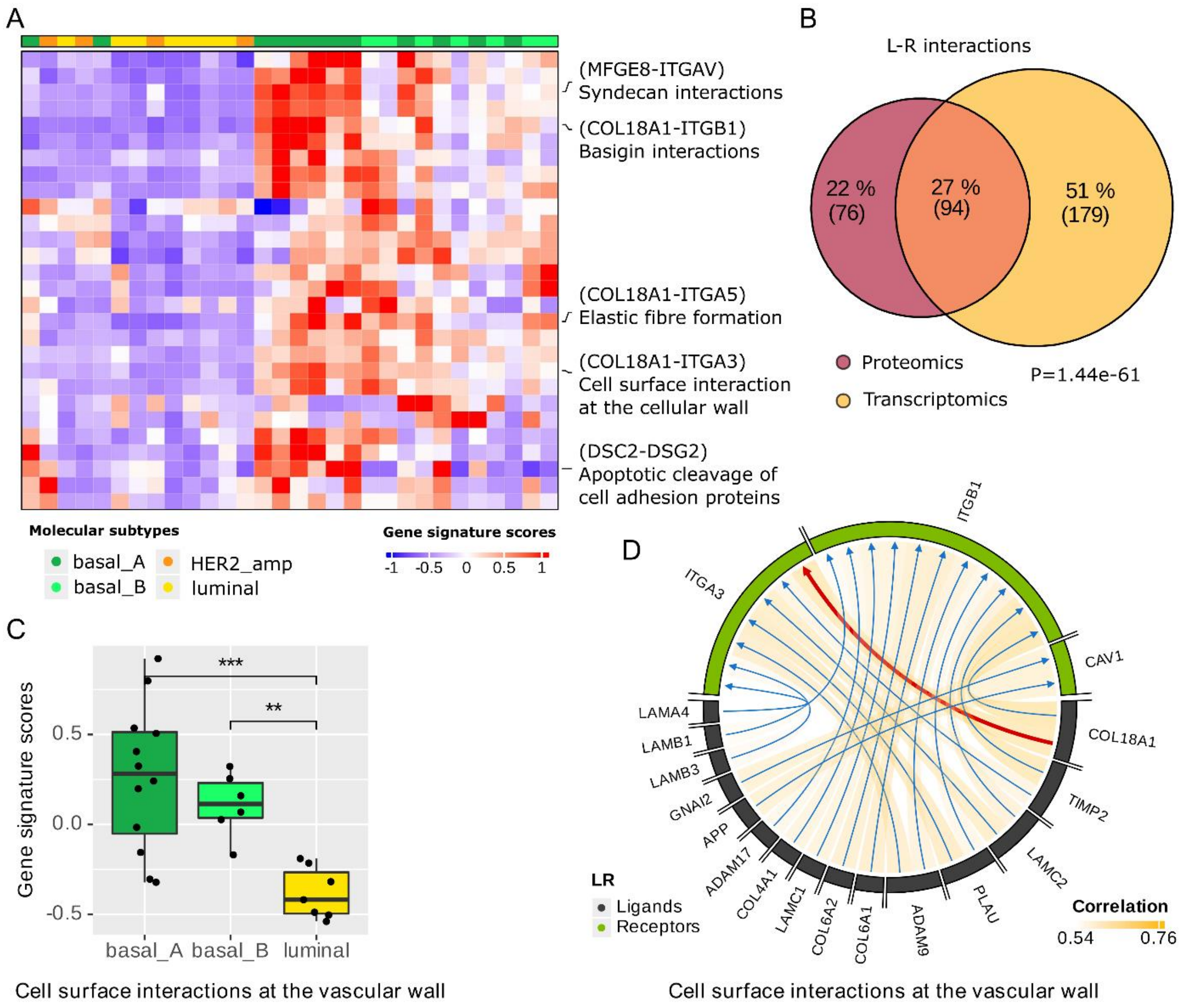


**Figure S12.** Analysis of breast cancer cell line proteomics data. (**A**) Gene signature scores at the pathway level. (**B**) Intersection of L-R interactions found in transcriptomics and proteomics. (Hypergeometric test.) (**C**) Boxplot showing differences in gene (protein here!) signature scores for the molecular subtypes (Wilcoxon test, ** P < 0.01, *** P < 0.001). (**D**) Chord diagram linking the most correlated ligands to their receptors in the Cell surface interactions at the vascular wall pathway. Highest correlation with a red arrow.

### Ortholog mapping: an illustration with a murine body map

To demonstrate BulkSignalR ability to work with nonhuman data we employed a transcriptomic murine body map covering 13 nonsexual tissues (27) (Supplementary Methods). The few lines of R code where a nonhuman species must be specified are illustrated in Figure S13. We note that working with a different species is essentially transparent.

```
# mouse expression data
m.counts <- …

# create the ortholog mapping
ortholog.dict <- findOrthoGenes(from_organism="mmusculus", from_values=rownames(m.counts))
h.counts <- convertToHuman(counts=m.counts, dictionnary=ortholog.dict)

# BulkSignalR Data Model
bsrdm <- prepareDataset(h.counts, species="mmusculus", conversion.dict=ortholog.dict)

# Inferences
bsrinf <- initialInference(bsrdm)
bsrinf <- resetToInitialOrganism(bsrinf, conversion.dict=ortholog.dict)

# the rest of the code is unchanged (reductions, gene signatures, etc.), refer to Figure S6
```

**Figure S13.** Code example working with nonhuman species.

We started the analysis at the pathway level for the sake of simplicity (Figure S14, Table S2). Reference to specific L-R interactions were obtained searching BulkSignalR output table, but also representing individual L-R interactions in a second heatmap (Figure S15).

Hierarchical clustering of gene signature scores at the pathway level grouped replicates of the same tissues (Figure S14). This indicated that differences in transcriptional programs between tissues related to different cellular networks. As reported by the data authors (27), the brain displayed the largest differences in its expression profiles compared to the other tissues. Naturally, high gene signature scores were associated with neurotransmitter signaling, neuronal migration and non-tyrosine kinases (nTRK) activity. The well-known interaction between the synaptic protein Nrxn2 (neurexin 2) and its binding partner Nxph3, which is a member of the trans-synaptic family of proteins called neurexophilins (28) featured the strongest gene signature score in the brain.

In contrast, the liver was characterized by the specific activation of plasma lipoprotein assembly, regulation of insulin-like growth factor and iron uptake and transport, processes known to be regulated by this organ (29–31). For instance, the regulation of Insulin-like Growth Factor (IGF) transport and uptake by Insulin-like Growth Factor Binding Proteins (IGFBPs) appears to be highly specific to the liver in Figure S14, in agreement with a liver contribution of roughly 75% of circulating IGF-I (32). A L-R interaction that was prevalent in the liver and the small intestine was Apob-Mttp (Figure S15). This interaction drives plasma lipoprotein assembly in the liver. In the intestine, Mttp facilitates the transfer of dietary and endogenous fat by assisting in the assembly and secretion of triglyceride-rich apolipoprotein B-containing lipoproteins (33). Of note, human lipoproteins are predominantly produced by the small intestine, explaining why this pathway was not detected in the large intestine (34). The interaction B2m-Hfe was also strong in the liver and expressed at a lower level in the spleen. This interaction plays a central role in iron transport and metabolism, which explains its tissue-specific localization (29, 35–38).

Spleen and thymus expression profiles harbored common traits and were very different from those of the other organs. These close transcriptomes were characterized by an expected upregulation of pathways associated with immune response and a downregulation of signaling processes related to tissue connectivity, *e.g.*, collagen formation. The Btla-Cd79a interaction was strongly upregulated in the spleen and the thymus (Figure S15). It is known to be related to BCR regulation in the spleen (39) and an increased expression of Btla through B cell maturation, as well as its specific association with Cd79a in the regulation of B cells, has also been described (40, 41). Btla can be extensively expressed in lymph nodes (absent from the body map), thymus and spleen, but little or no expression has been detected in other organs (42) as it was the case in our analysis.

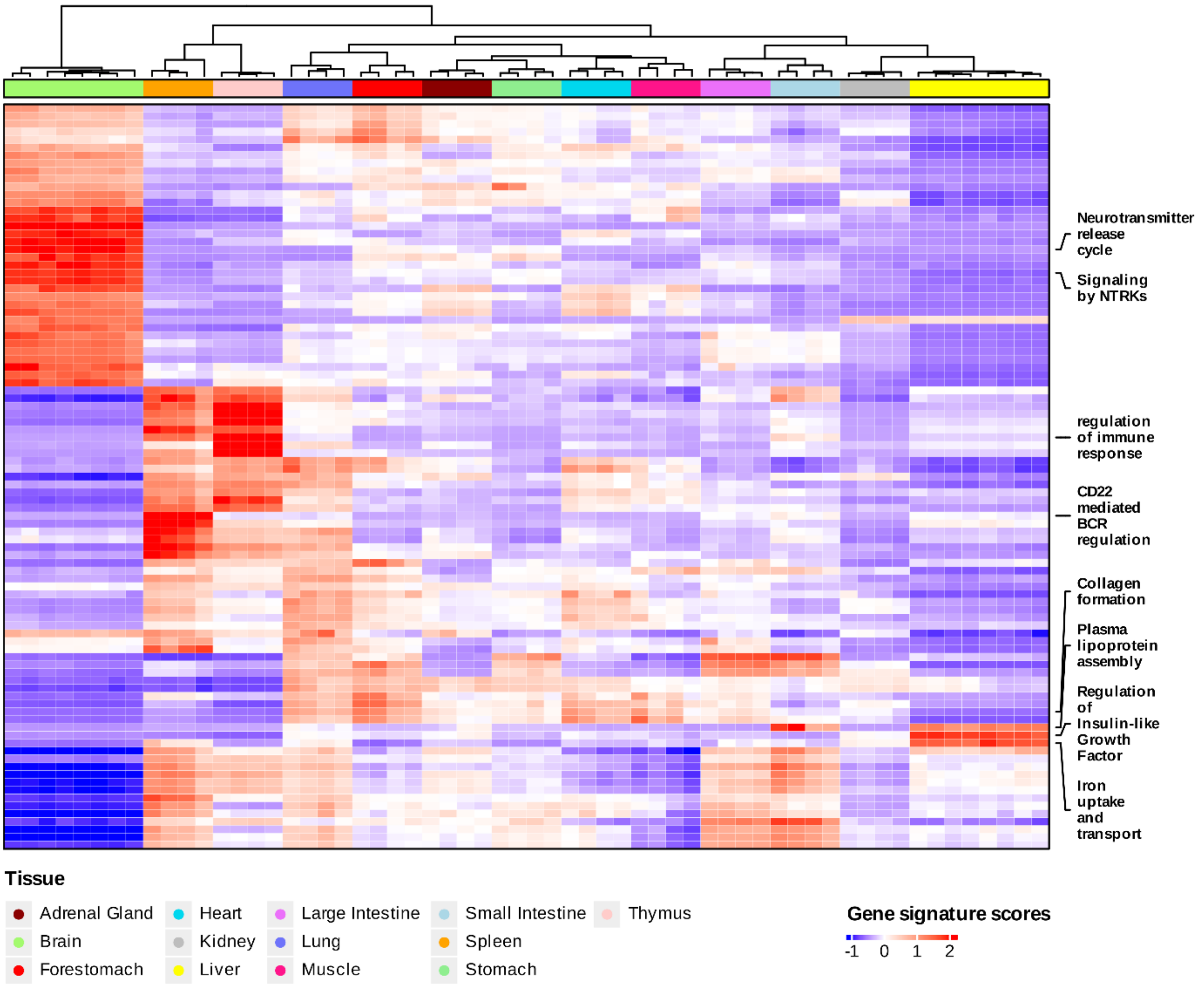


**Figure S14.** BulkSignalR analysis reduced at the pathway level for mouse transcriptomes (27).

From the unique L-R interaction heatmap (Figure S15), we can read that the gene signature score of the Desmocollin-2 (Dsc2)-Desmoglein 2 (Dsg2) interaction, which are the major desmosomal cadherins of digestive epithelia (43), was stronger in the intestine (large and small) as well as in the stomach and forestomach. Also from Figure S15, we see that several L-R interactions can be associated with diverse tissues more or less specifically. For example, the Col4a5-Itga1 interaction is indicative of the presence of smooth muscle fibers, which are located in walls of hollow visceral organs, the cardiovascular system and the lungs. Indeed, the gene signature scores for this interaction was predominantly higher in lung, heart, and forestomach, and moderate in the intestine, muscles, stomach, adrenal gland, and kidney.

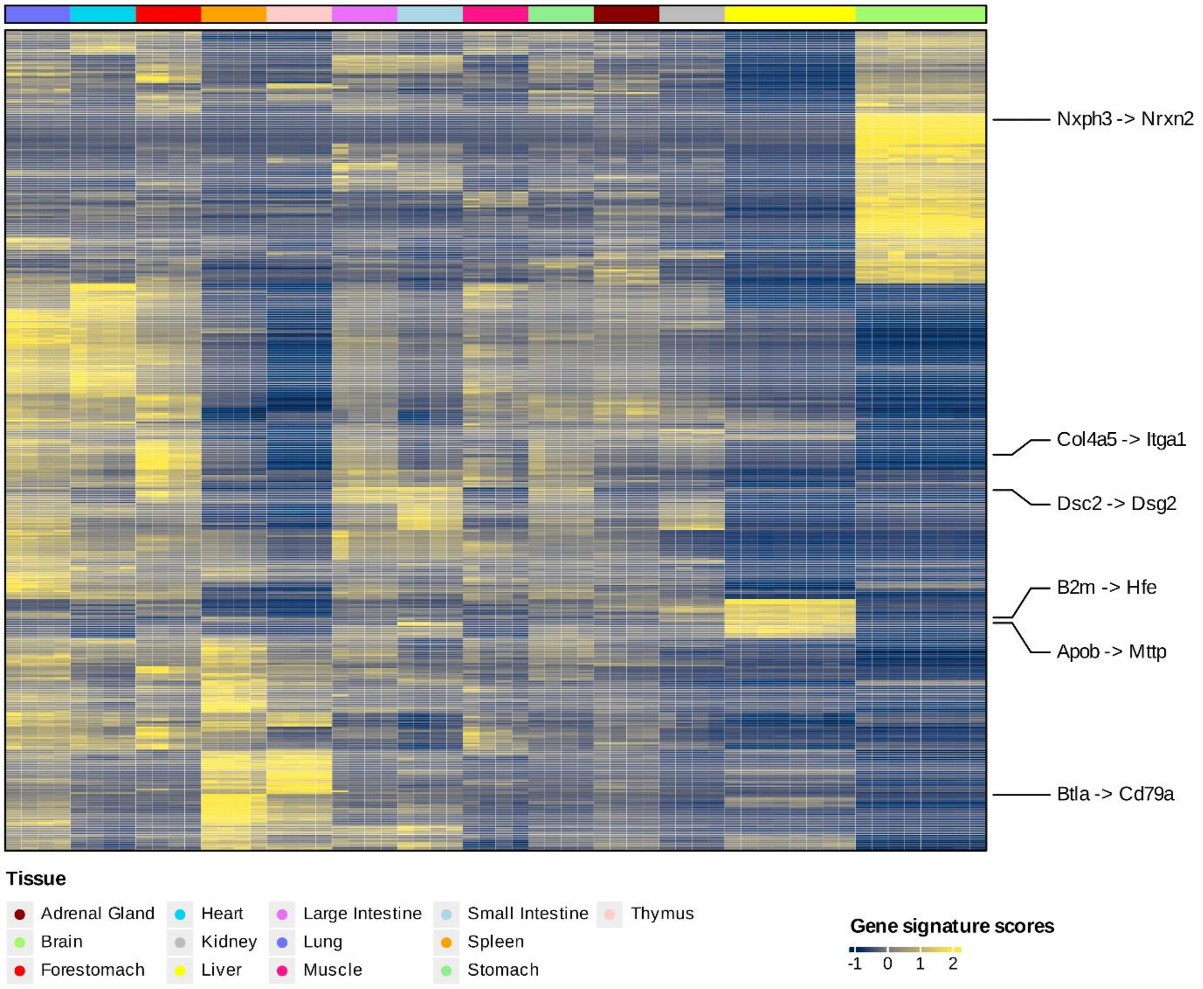


**Figure S15.** BulkSignalR analysis reduced to unique L-R interactions for mouse transcriptomes (27)

```
# read expression data
counts <- read.csv(...)
# or get them from a Seurat object
counts <- as.matrix(seurat.object@assays$Spatial@counts)

# standard pipeline with adapted parameters
bsrdm <- prepareDataset(counts, min.count=1, prop=0.01)
bsrdm <- learnParameters(bsrdm, min.positive=2)
bsrinf <- initialInference(bsrdm , min.cor=-1)

# signature scores, check correlation between L and R is positive because of min.cor=-1 above
bsrinf.red <- reduceToBestPathway(bsrinf)
pairs.red <- LRinter(bsrinf.red)
s.red <- getLRGeneSignatures(bsrinf.red, qval.thres=1)
scores.red <- scoreLRGeneSignatures(bsrdm, s.red) [pairs.red$LR.corr>0.02 & pairs.red$qval<0.01,]

# get spatial coordinates and tissue areas (if available), those data must be in a data.frame that we
# call "areas" for this example script
# idSpatial array_col array_row ground_truth
# 50x102       102        50       Label3
#   3x43        43         3       Label1
#  59x19        19        59       Label3

# spatial plot, for instance the 20th interaction
title <- gsub("\\}", "", gsub("\\{", "", rownames(scores.red)[20]))
spatialPlot(scores.red[20,], areas, title)

# generate visual index
spatialIndexPlot(scores.red, areas, "file-name.pdf")

# statistical association with tissue areas
assoc.bsr <- spatialAssociation(scores.red, areas)
spatialAssociationPlot(assoc.bsr)

# 2D-projection of score spatial distribution patterns
spatialDiversityPlot(scores.red, assoc.bsr, with.names=TRUE)
```

**Figure S16.** Spatial data pipeline.

### A spatial triple-negative breast cancer dataset

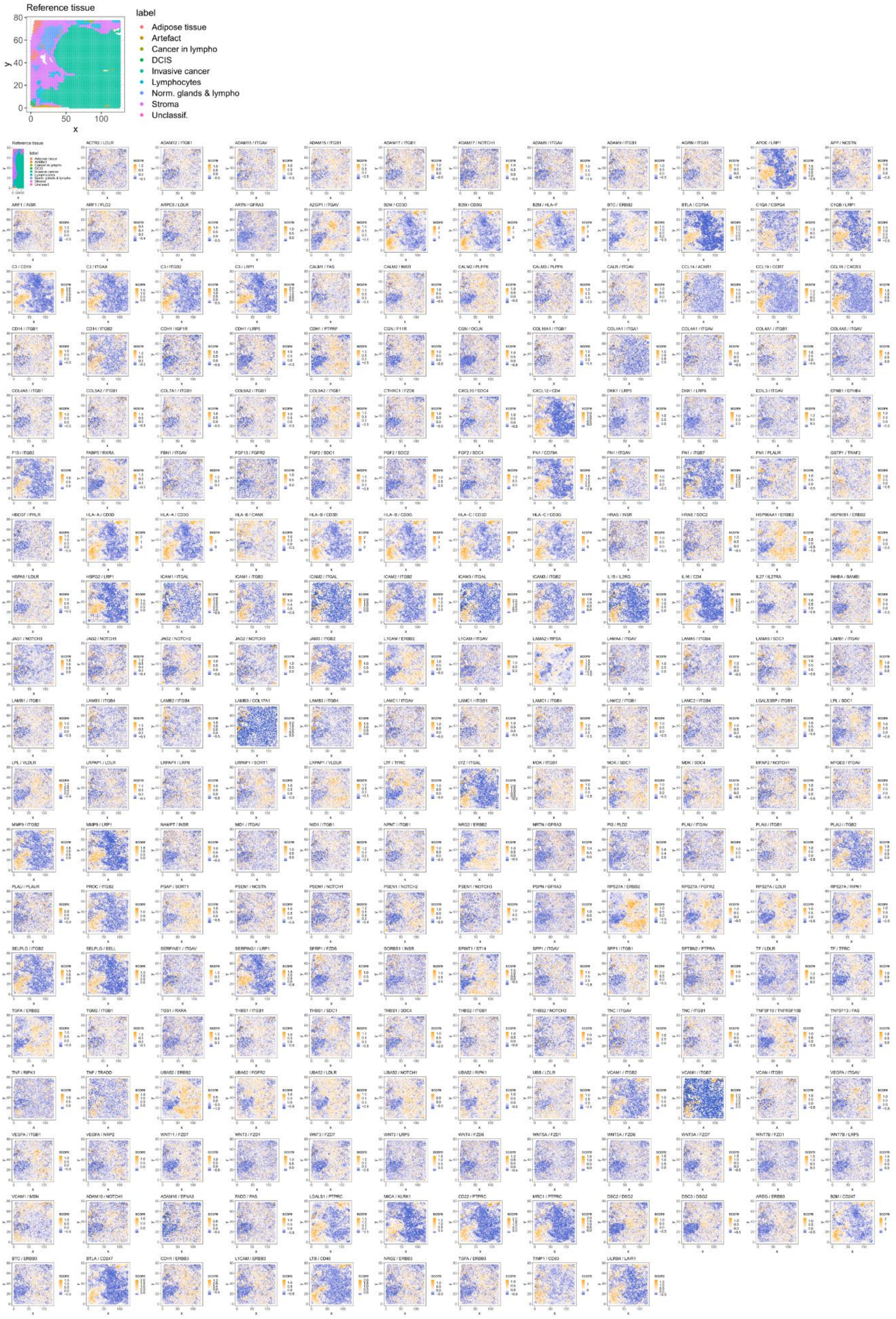

**Figure S17.** Overview of the 224 identified interactions in the TNBC sample (44) represented by the spatial rendering of their gene signature score.

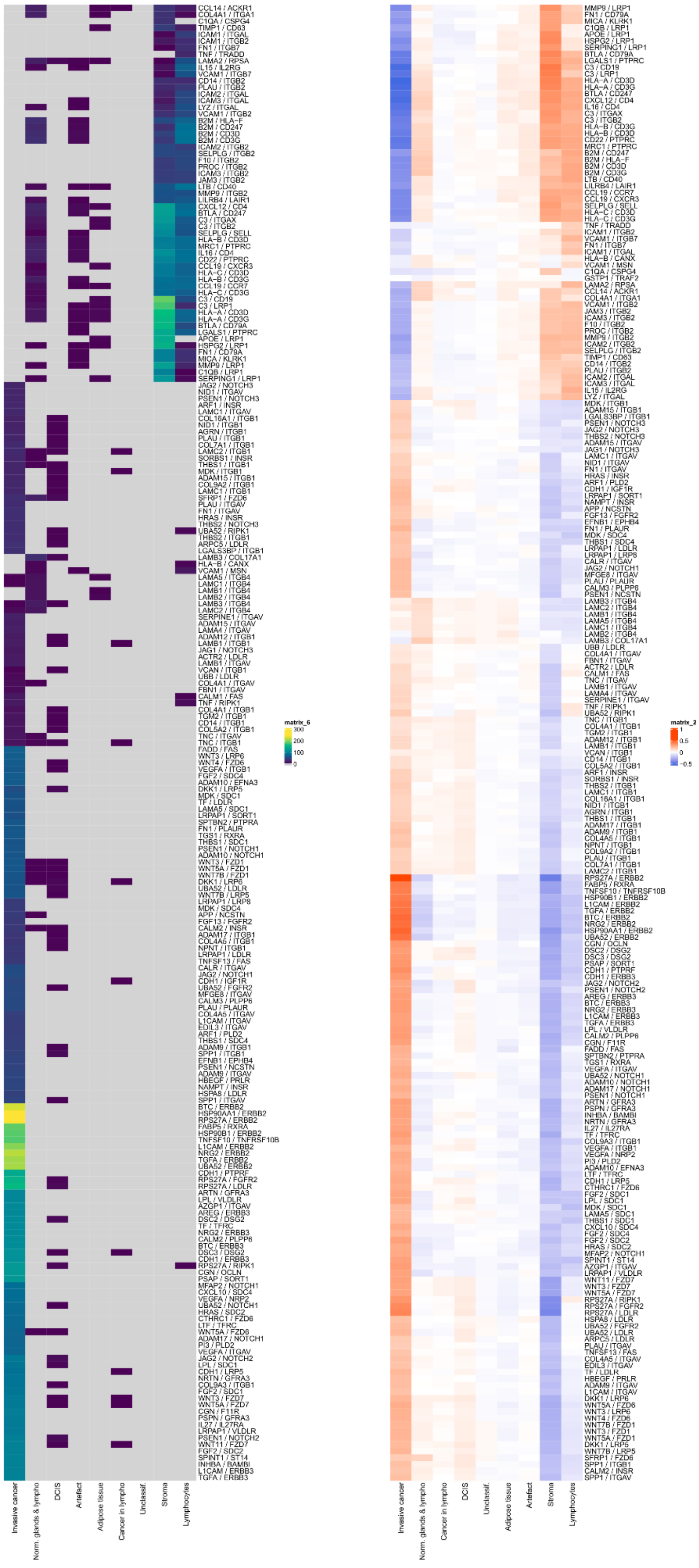

**Supplementary Figure 18.** Spatial associations in the TNBC dataset (left Wilcoxon test adjusted P-values, right Spearman correlations).

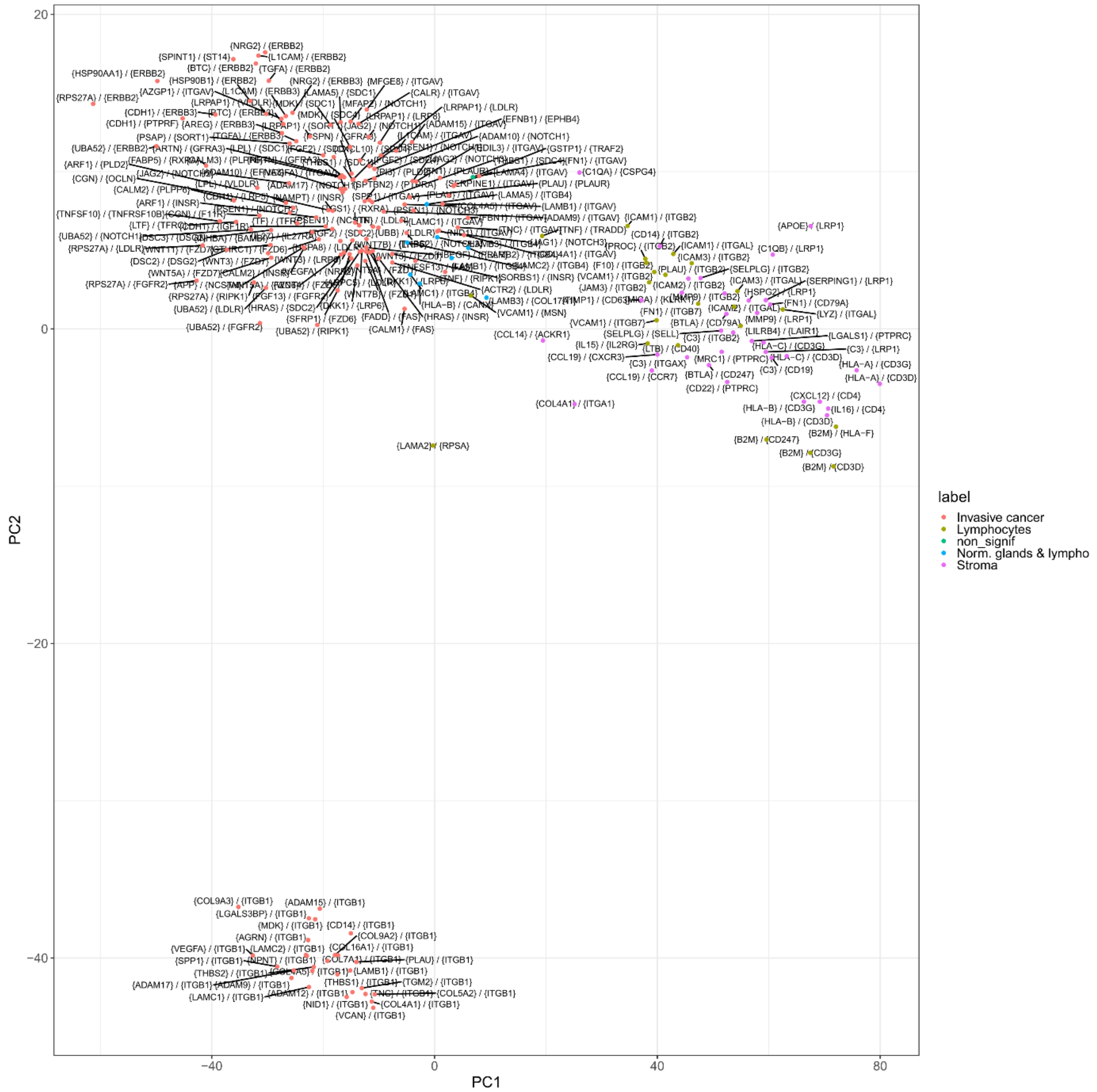


**Supplementary figure 19.** Two-dimensional projection (PCA) of L-R interaction gene signature score spatial patterns (shown in Figure S17). Dot colors represent the strongest spatial area association (according to Figure S18). Projections using t-SNE are available too.

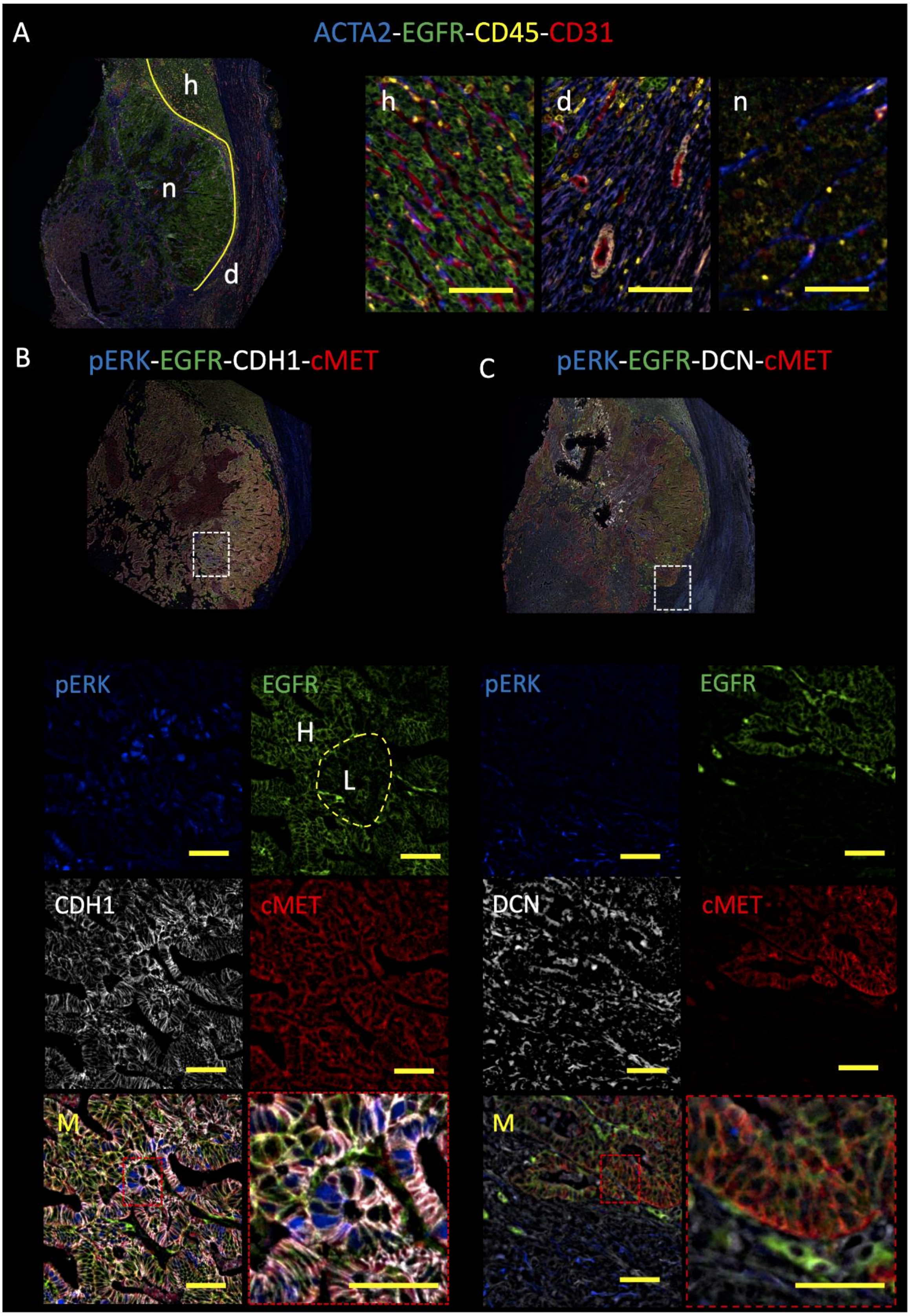


**Figure S20. Figure 6.** Immunofluorescence validation of selected ligand-receptor interactions in CRCLM2. (**A**) (*left*) Structural overview of CRCLM1 with stromal area (delineated in white) as well as green labelled cancer cells. ACTA2 is CAF marker, CD45 marks immune cells, EGFR cancer cells and CD31 labels endothelial cells. Area marked (d) denotes desmoplastic reaction while (n) denotes necrosis. (*right*) High-power view of desmoplastic and necrotic areas. (**B**) Interaction analysis of CDH1-EGFR and CDH1-cMET. Low-power view (100x) of quadruple staining pERK, EGFR, CDH1, and cMET in CRCLM2. The area of interest is highlighted and further shown in panels below. In the EGFR panel, two zones can be delineated corresponding to high (H) and low (L) receptor levels. (**C**) Same as in (B) for the DCN-EGFR and DCN-cMET interactions.

### Supplementary Methods

#### Bulk datasets and their preparation

The Genotype-Tissue Expression (GTEx) Project provided us with brain frontal cortex, liver, and pancreas transcriptomes. These RNA-seq data were obtained from v8 (2017-06-05) RNASeQCv1.1.9_gene_reads_gct. SDC data are available from GEO (GSE138581) and from Ref. (2) authors. TCGA HNSC transcriptomes were downloaded (gene read counts) from the BROAD Institute GDAC at firebrowse.org. These RNA-seq data were analyzed with BulkSignalR default parameters.

In Figure S1, we reported a ROC curve based on sarcoma microarray data(3) (Affymetrix). The data were downloaded from GEO (GSE21050). BulkSignalR default parameters were used. We also used a functionality of BulkSignalR data preparation function, which is to eliminate redundant rows in the expression matrix (for the same gene) due here to redundancy in Affymetrix probes. The row with maximal sum across all the samples was chosen.

In Figure S9 and following, we analyzed breast cancer cell line transcriptomes (RNA-seq) and proteomes. Data were retrieved from DepMap (12) 22Q2 Portal and processed with BulkSignalR default parameters except for transcriptomics where min.count was set to 1 and log.transformed to TRUE. The breast cancer cell line dataset null model was forced to censored normal (instead of Gaussian kernel-based empirical) to be the same as for the lung (main paper Figure 2) that was automatically set as censored normal by our algorithm.

Mouse body map transcriptomes(27) were processed with default parameters, but min.count that was set to 1. BulkSignalR ortholog mapping mechanism was used to convert murine genes to human and back.

In every case, the FDR threshold for selecting interactions in bulk was 0.1% and 1% in spatial trabscriptomics.

#### Single-cell datasets

In Figure S4, we provide a comparison of the number of L-R interactions found by BulkSignalR from three bulk datasets with SingleCellSignalR (45) results from as similar as possible tissues in single-cell. The single-cell datasets used were pancreas (4), liver (5, 6), and brain cortex (7, 8). Data were prepared with SingleCellSignalR default pipeline and parameters. Namely, each dataset was normalized and filtered (non-expressed genes) with the function `data_prepare()` (default parameters), the cell populations were found by calling `clustering()` (default parameters, calls SIMLR (46)), and the L-R interactions, both autocrine and paracrine, we found by calling `cell_signaling()` (default parameters, but LRscore threshold set at 0 to filter *a posteriori*). Finally, in the comparison with BulkSignalR output, L-R interactions found in the single-cell data were considered correct provided their LRscore was above a conservative threshold of 0.6 (45).
